# Complementary *In Vivo* Approaches Reveal Quiescent Melanocytes in Anagen Outside the Hair Follicle Bulge

**DOI:** 10.1101/2020.05.27.117051

**Authors:** Bishal Tandukar, Sandeep S. Joshi, Li Pan, Thomas J. Hornyak

## Abstract

Melanocyte stem cells (McSCs) are key components of the hair follicle (HF) stem cell system that are derived from neural crest during embryogenesis and are responsible for regeneration of differentiated melanocytes during successive HF cycles. Our previous research has shown presence of two subsets of phenotypically and functionally distinct McSCs exist in murine telogen HFs, CD34+ McSCs in the bulge/lower permanent portion (LPP) and CD34− McSCs in the secondary hair germ (SHG). Whether these subsets are maintained independently or exist in a developmental hierarchy is not yet known. Using *Dct*-H2BGFP mice, we analyzed the quiescent and proliferative properties of McSCs and melanocytes in anagen and telogen. We found unexpectedly that Kit+Nestin− quiescent melanocytes are maintained outside of the bulge/LPP region throughout anagen in addition to the Kit+Nestin+ quiescent melanocytes of the bulge/LPP. Both subpopulations express lower levels of melanocyte differentiation markers Mitf, Pax3, Dct, Tyrp1 and Tyr compared to differentiated melanocytes of the HF bulb/matrix. These results suggest that quiescent melanocytes localized in the outer root sheath, both in and below the bulge/LPP) retain the stem cell phenotype observed in quiescent McSCs during telogen. This finding has implications for maintenance of distinct subsets of McSCs throughout successive HF cycles.

## INTRODUCTION

Melanocyte stem cells (McSCs) are quiescent melanocytic progenitors present in the bulge and subbulge regions of the telogen hair follicle (HF), expressing *Dopachrome tautomerase (Dct)* and retaining a bromodeoxyuridine (BrdU) label administered during early anagen (Nishimura et al. 2002). Notch signaling, acting through RBP-J (Hamaguchi et al. 1989) and Hes1 (Kageyama et al. 2000), facilitates survival of McSCs and their precursors during development (Moriyama et al. 2006). Collagen XVII expressed by epithelial HF stem cells (HFSCs) induces a TGF-β signal from HFSCs to McSCs inhibiting premature melanocyte differentiation, thereby promoting McSC maintenance (Nishimura et al. 2010; Tanimura et al. 2011). A Wnt signal from HFSCs in the secondary hair germ (SHG) (Ito et al. 2004) of the telogen HF activates β-catenin upon anagen onset, stimulating melanocyte proliferation and differentiation coordinated with HFSC activation and growth. Finally, the membrane protein CD34 distinguishes CD34+ bulge McSCs from CD34− SHG McSCs during HF telogen, with these subsets exhibiting distinct functional properties (Joshi et al. 2019).

Epithelial HFSCs have been extensively studied, establishing the paradigm of the HF bulge (Cotsarelis et al. 1990) and SHG (Ito et al. 2004) as quiescent niches. The lateral disc hypothesis was proposed to explain the origin of SHG epithelial stem cells and their distinct identity from bulge stem cells. The hypothesis posited that a group of *Shh-*expressing cells present within the anagen matrix contributes to the SHG during telogen (Panteleyev et al. 2001). However, this hypothesis was refuted subsequently after *ShhCre*^*ER*^*/Rosa-LSL-LacZ* mice were used to trace these cells or their progeny from late anagen to a distinct set of inner root sheath cells in telogen, revealing the presence of slow-cycling cells early in the retracting epithelial strand and later in both the SHG and bulge. Although the interpretation of the results of these and related experiments was that SHG cells are derived directly or indirectly from the bulge (Greco et al. 2009), these data may also support a model whereby SHG and bulge cells are maintained quasi-independently. No similar studies have been reported to date to examine specifically the relationship between quiescence and proliferation of McSCs located in the bulge and SHG.

In this study, we dissect proliferative differences between McSCs located in the HF bulge and other HF regions during both telogen and anagen. We utilize BrdU incorporation studies in conjunction with a bitransgenic mouse strain, *Dct-*H2BGFP (Joshi et al. 2019; Joshi et al. 2018), for identification of McSCs. We reveal the unexpected presence of a quiescent population of melanocytes residing outside the bulge region during HF anagen. Using doxycycline regulation of *Dct-*H2BGFP mice, we confirm these observations and demonstrate how dilution of the GFP label correlates with melanocyte proliferative status. Marker analysis of McSCs and melanocytes in the HF bulge, bulb, and regions in between reveals even greater heterogeneity of HF melanocytes.

## RESULTS

### Quiescence of McSCs throughout telogen

Prior characterization of McSCs using *in vivo* BrdU labeling determined quiescence through their retention of BrdU incorporated during prior anagen. This analysis (Nishimura et al. 2002) identified cells quiescent at a single time point during HF telogen. It did not explore whether most or all HF McSCs remained quiescent throughout telogen. Similar studies conducted in HFSCs (Greco et al. 2009) were focused upon the epithelial HF stem cell component and did not use a specific marker to determine the proliferative or quiescent status of McSCs throughout telogen.

To characterize comprehensively quiescence of McSCs throughout telogen, *Dct*-H2BGFP mice (Joshi et al. 2018) were intraperitoneally injected with BrdU every 12 hours throughout early (P49-P56), mid-(P56-P63), and late telogen (P63-P70) (Figure 1a). BrdU incorporation was determined for bulge or SHG McSCs respectively. Of telogen-stage follicles analyzed, results showed that 98.0±1.3% of SHG McSCs and 97.4±1.3% of bulge McSCs were BrdU-during P49-P56, 97.7±0.9% of SHG McSCs and 97.2±0.5% of bulge McSCs were BrdU-during P56-P63, and 96.5±1.8% of SHG McSCs and 98.9±1.1% of bulge McSCs were BrdU-during P63-P70 (Figures 1b,c). Compared to vehicle-injected mice, BrdU incorporation signal was not significant, suggesting that essentially all HF McSCs are quiescent throughout telogen. The results support the conclusion that bulge and SHG McSCs are each quiescent populations throughout telogen and do not proliferate to repopulate either population during this stage.

**Figure 1.**
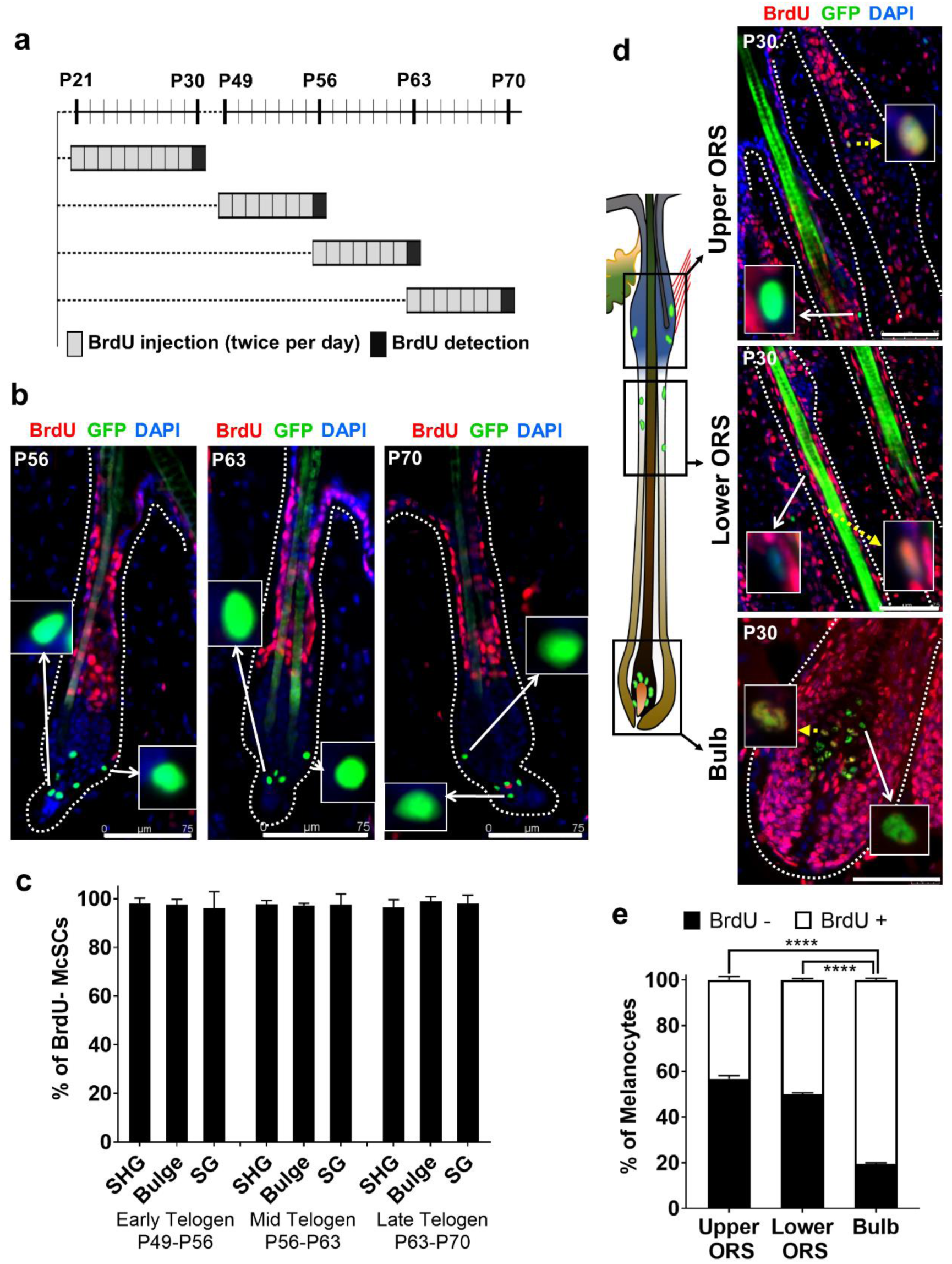
Characterization of proliferating and quiescent follicular melanocytes during anagen onset and telogen. (a) Experimental design depicts BrdU administration during anagen onset (P21-P30), and early (P49-P56), mid (P56-P63) and late (P63-P70) telogen. (b) Analysis of BrdU incorporation during telogen stages. Inset images show absence of BrdU incorporation in bulge and SHG McSCs. (c) Quantification of BrdU incorporation in SHG, bulge and SG McSCs at different telogen stages. (mean ± SEM, *n* = 3, *P* >0.1 for all 3 telogen stages vs control) (d) BrdU incorporation in upper ORS/bulge, lower ORS and bulb during anagen onset (P21-P30). Inset images show BrdU+ (yellow dotted arrows) and BrdU-(white arrows) melanocytes. (e) Quantification of BrdU+ and BrdU-melanocytes show quiescent melanocytes in both upper ORS and lower ORS compared to bulb (mean ± SEM, *n* = 3). \*\*\**P* < 0.0001 (one-way ANOVA). Scale bars, 75µm. See also figure S2 and table S1.

Moreover, 34% of HFs analyzed at P70 had entered anagen, consistent with the enhanced uptake of BrdU in the SHG noted previously (Greco et al. 2009). The incorporation of BrdU administered from 7 days prior in three distinct regions of the anagen follicle, the bulge/upper outer root sheath (ORS); the lower ORS, here defined as the region between the bulge and the bulb; and the bulb were quantified. Of the anagen follicles, 18.6±4.2%, 30.3±1.1% and 74.1±2.2% of Dct-expressing cells of the bulge/upper ORS, lower ORS, and bulb respectively were BrdU-positive, compared to telogen follicles at P70 and prior in telogen, indicating transition into the proliferative stage of the HF cycle (Figure S1b,c). The exact stage of the HF cycle was confirmed by alkaline phosphatase staining (Handjiski et al. 1994; Paus et al. 1999; Tobin et al. 1999) (Figure S1a). A small number of GFP+ McSCs in early, mid and late telogen were observed located above the bulge closer to the sebaceous gland in the BrdU+ infundibular region of the HF (Figure S2). They were BrdU- (Figure S2a, Table S1), despite being surrounded by GFP− BrdU+ cells in this proliferating zone and were not explored further.

### Quiescence and proliferation of melanocytes and McSCs during anagen

McSCs that are dihydroxyphenylalanine-stained and negative (DOPA−), indicating an undetectable level of tyrosinase activity, in all HF cycle stages are maintained in the outer root sheath (ORS) layer of the HF, unlike DOPA+, differentiated melanocytes located within the matrix of the anagen hair bulb that pigment the hair shaft (Tobin et al. 1999). Preliminary observations showed GFP− expressing melanocytes in the ORS between the bulge and the matrix, usually limited to the upper half of the anagen HF (Figure 1d). We define the location of these cells as the lower ORS, as they are outside of the bulge/lower permanent portion (LPP) which we refer to as the upper ORS during anagen. The upper ORS/bulge and lower ORS regions described here correspond closely to the LPP and the upper transient portion (UTP) previously defined (Harris et al. 2013). However, this study was performed using depilation-induced anagen, rather than endogenous anagen, and used the junction of the dermis and subcutis, not intrinsic follicular structures, to define the boundary between LPP/UTP. In first anagen, we recognize the bulge/upper ORS as an epithelial swelling around the club hair (Cotsarelis et al. 1990; Myung and Ito 2012) corresponding to the region of positive expression of CD34 (Figure S4) (Joshi et al. 2019). We sought to define whether lower ORS follicular melanocytes during anagen are quiescent or proliferative.

To determine the proliferative state of ORS melanocytes, BrdU labelling was performed *in vivo* during anagen onset from P21-P30 to identify cells that had proliferated (Figure 1a). Analysis showed that a substantial fraction of Dct-expressing cells in the upper ORS, lower ORS, and bulb of anagen HFs (56.5±1.5%, 49.9±0.6% and 19.4±0.7%) featured no BrdU label (Figure 1d,e). These results indicate that a quiescent population of melanocytes is maintained throughout early anagen in all regions of the HF.

We were surprised that a substantial proportion of quiescent melanocytes at P30 could be identified despite the extensive cellular proliferation associated with follicular anagen. To confirm this finding, we used doxycycline (Dox) regulation of *Dct*-H2BGFP mice to extinguish ongoing transgene expression, similar to the technique used previously to extinguish *Krt5*-driven expression in proliferating murine follicular keratinocytes. The residual level of GFP expression is inversely correlated to cell proliferation during the period of Dox administration (Tumbar et al. 2004). Doxycycline was administered from P19-P30. Mice were examined for retention or dilution of a stable H2BGFP label in follicular melanocytes (Figure 2a). In comparison to control mice (Figure 2b), fluorescence images from Dox-treated mice show lower levels of GFP expression in the HF bulb compared to the upper and lower ORS regions, where numerous high GFP+ cells were still visible (Figure 2c). Quantification (n=3, 100 HFs/mouse, Table S3c) revealed distinct differences in the retention of high GFP expression following Dox administration, with 63±5% of GFP+ cells in the bulge/upper ORS and 62±3% of total GFP+ cells in the lower ORS retaining high GFP expression which was shown by only 11±3.8% of melanocytes in the HF bulb (Figure 2c). Heterogeneity in GFP expression was examined in control mice not receiving Dox. Control mice (n=3, 100 HFs per mouse) showed that only 10±0.9% of melanocytes in the upper ORS, 14±1.9% in the lower ORS, and 6±1.5% in the bulge have low GFP based on the GFP intensity threshold (Figure S3a) for GFP^high^ and GFP^low^ classes. Overall, we observed 20±5% (1368±44 GFP+ cells/100 HFs in Dox-treated mice vs 1720±78 GFP+ cells/100 HFs in controls) fewer total GFP+ cells in Dox-treated mice compared to controls (n=3, p<0.01). This difference is likely attributable to complete loss of GFP expression below the threshold of detectability in rapidly-dividing cells. Closer examination showed that the reduction of GFP+ total cell numbers between Dox-treated and control mice was restricted to the HF bulb (Figure S3b, Table S3a,b).

**Figure 2.**
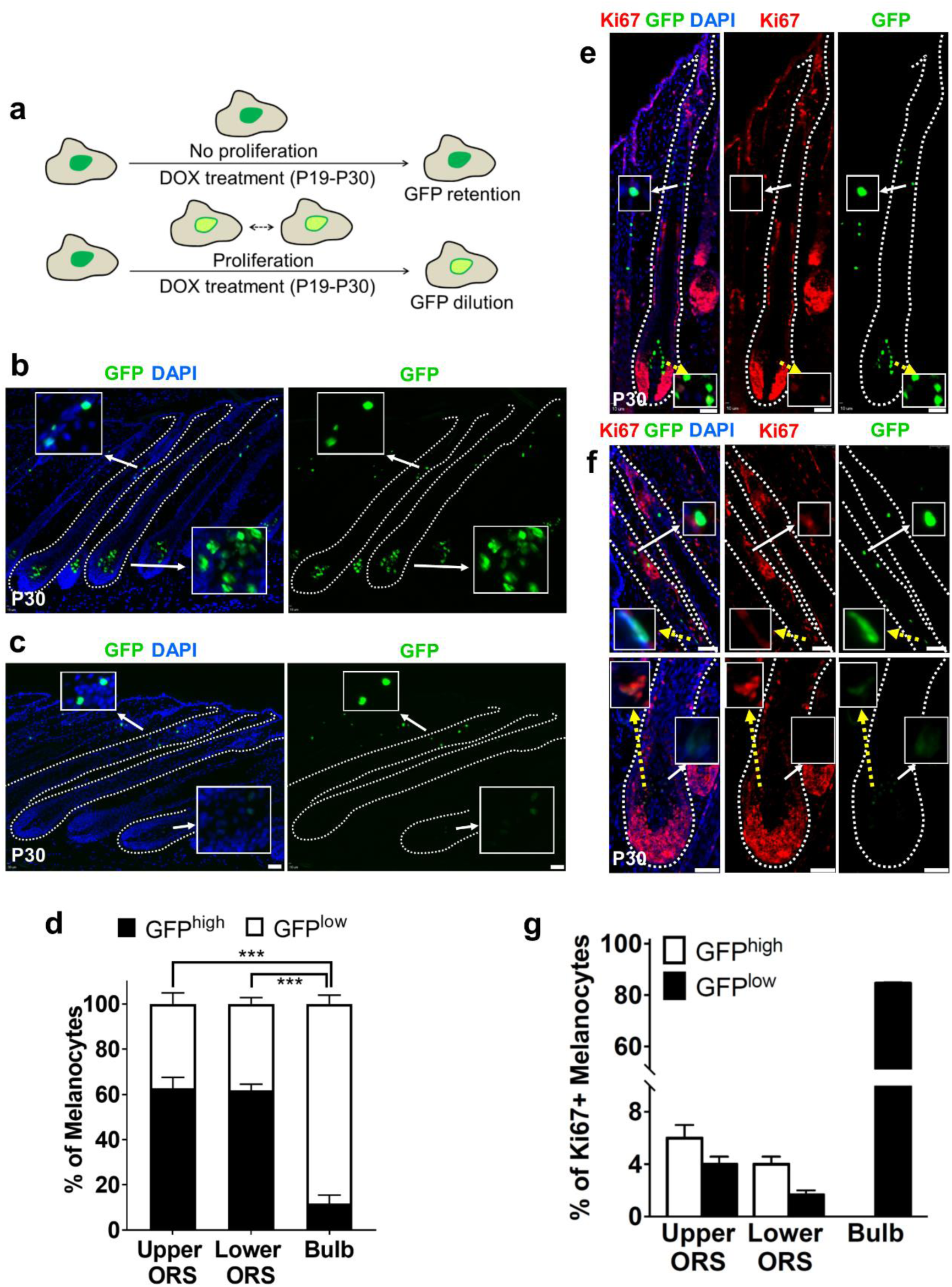
Characterization of proliferating melanocytes during anagen onset by doxycycline-induced GFP depletion. (a) Schematic representation of the model. GFP expression in melanocytes during anagen without (b) and with DOX treatment (c) from P19-P30. Inset images (c) show GFP^high^ (top) and GFP^low^ (bottom) melanocytes in ORS and bulb respectively. Scale bars, 50µm. (d) Quantification of GFP^high^ and GFP^low^ melanocytes after DOX treatment. (mean ± SEM, *n* = 3). \*\*\**P* < 0.001 (one-way ANOVA). (e) Ki67 expression in anagen HF (f) Ki67 expression in GFP^high^ cells from ORS (top panel) and GFP^low^ cells (bottom panel) from bulb after DOX treatment. Scale bars, 50µm. (g) Quantification of Ki67 expression in GFP^high^ and GFP^low^ melanocytes show correlation between GFP loss and Ki67 expression. (mean ± SEM, *n* = 3; 34 HFs/animal) χ^2^ (2 degrees of freedom, N = 664) = 414, \*\*\*\**P* < 0.0001 (Pearson’s chi-squared test). See also table S3.

Sox10 is expressed in McSCs where its expression at proper levels is critical for maintenance and repressing differentiation, and is less variable than other McSC markers (Harris et al. 2013). Consistent with these findings, we observed that Sox10 was expressed at similar levels in GFP+ melanocytes in the ORS and bulb regions (Figure 5a, Figure S13). Thus, we used Sox10 to identify melanocytes that may have lost detectable expression of GFP following administration of doxycycline. Comparison of co-localized expression of Sox10 and GFP in P30 anagen HFs between untreated and Dox-administered mice (Figures 5a,b; Figure S13) shows a selective absence of GFP in Sox10+ bulb melanocytes in Dox-administered mice. Quantification of immunofluorescence images showed significant numbers of Sox10+GFP− cells (80/376 Sox10+ cells, 21±3%) in the HF bulb in Dox-treated mice compared to controls (10/254 Sox10+ cells, 4±1%, p<0.001). In contrast, few or no Sox10+GFP−cells were observed in the ORS region where melanocytes reside (5/31 (14±5%) Sox10+ cells in upper ORS; 0/41(0%) Sox10+ cells in lower ORS region in Dox-treated mice versus 0/36 (0%) Sox10+ cells in upper ORS; 0/47(0%) Sox10+ cells in lower ORS region of control mice) (Figure S3c, Table S3e). The selective absence of GFP+Sox10+ melanocytes in the HF bulb of Dox-treated mice suggests that the most highly proliferative melanocytes are restricted to this location during anagen. Persistence of detectable GFP expression in the lower ORS is consistent with our finding (Figure 1d,e) of quiescent melanocytes existing outside of the bulge region during HF anagen.

**Figure 3.**
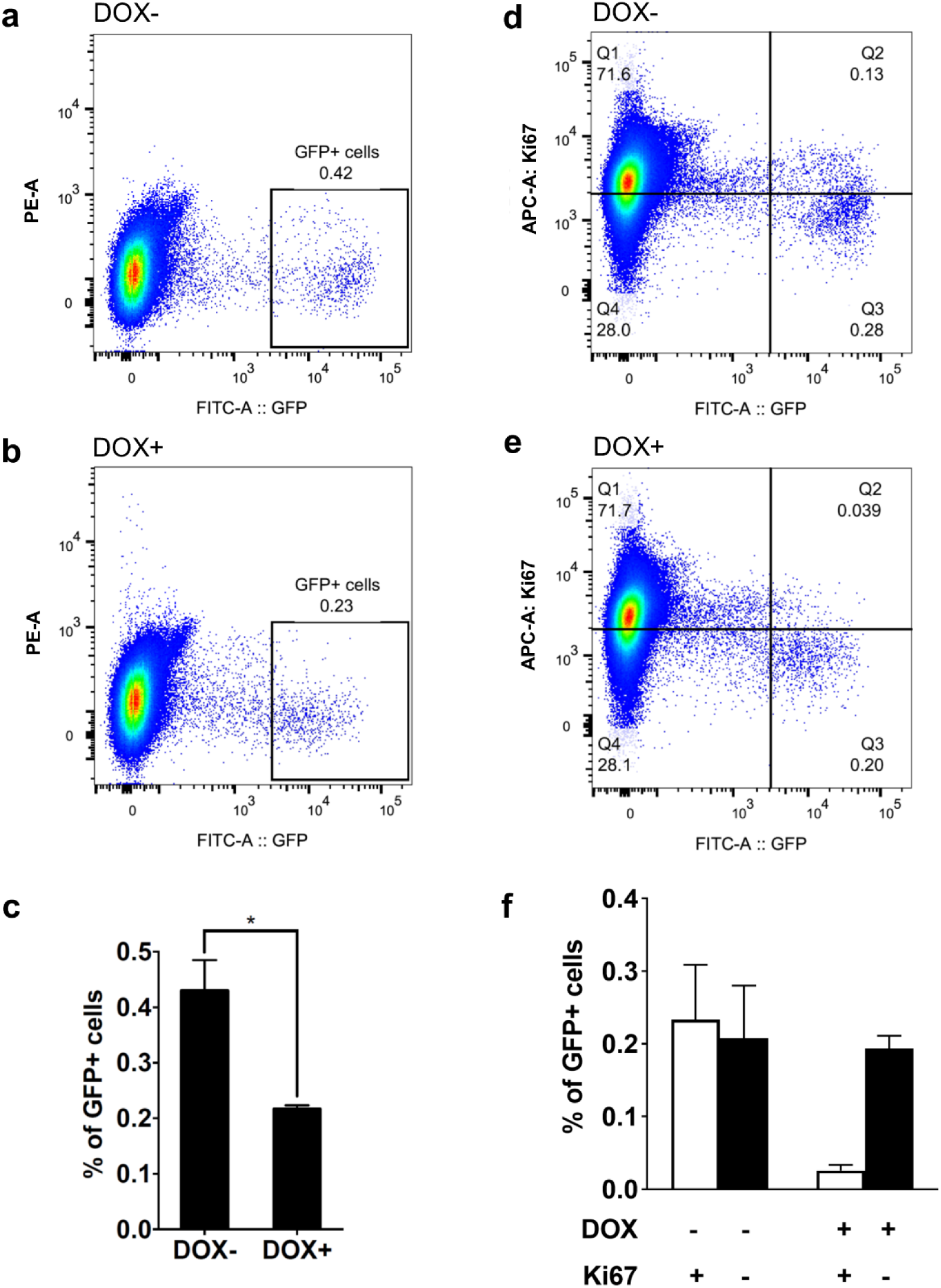
Correlation between GFP loss and Ki67 expression in melanocytes after DOX treatment. (a) Flow cytometry analysis of GFP+ cells with (a) and without (b) DOX treatment. (c) Quantification of total GFP+ cells from mice +/- DOX. (mean ± SEM, *n* = 3). \**P* < 0.05 (Student’s t-test). Two-color flow cytometry analysis of GFP and Ki67 expression without (d) and with (e) DOX treatment. (f) Quantification of Ki67+ and Ki67-melanocytes identified +/- DOX treatment. (mean ± SEM, *n* = 3) *P* = 0.05 Ki67+ vs Ki67- melanocytes (Student’s t-test).

**Figure 4.**
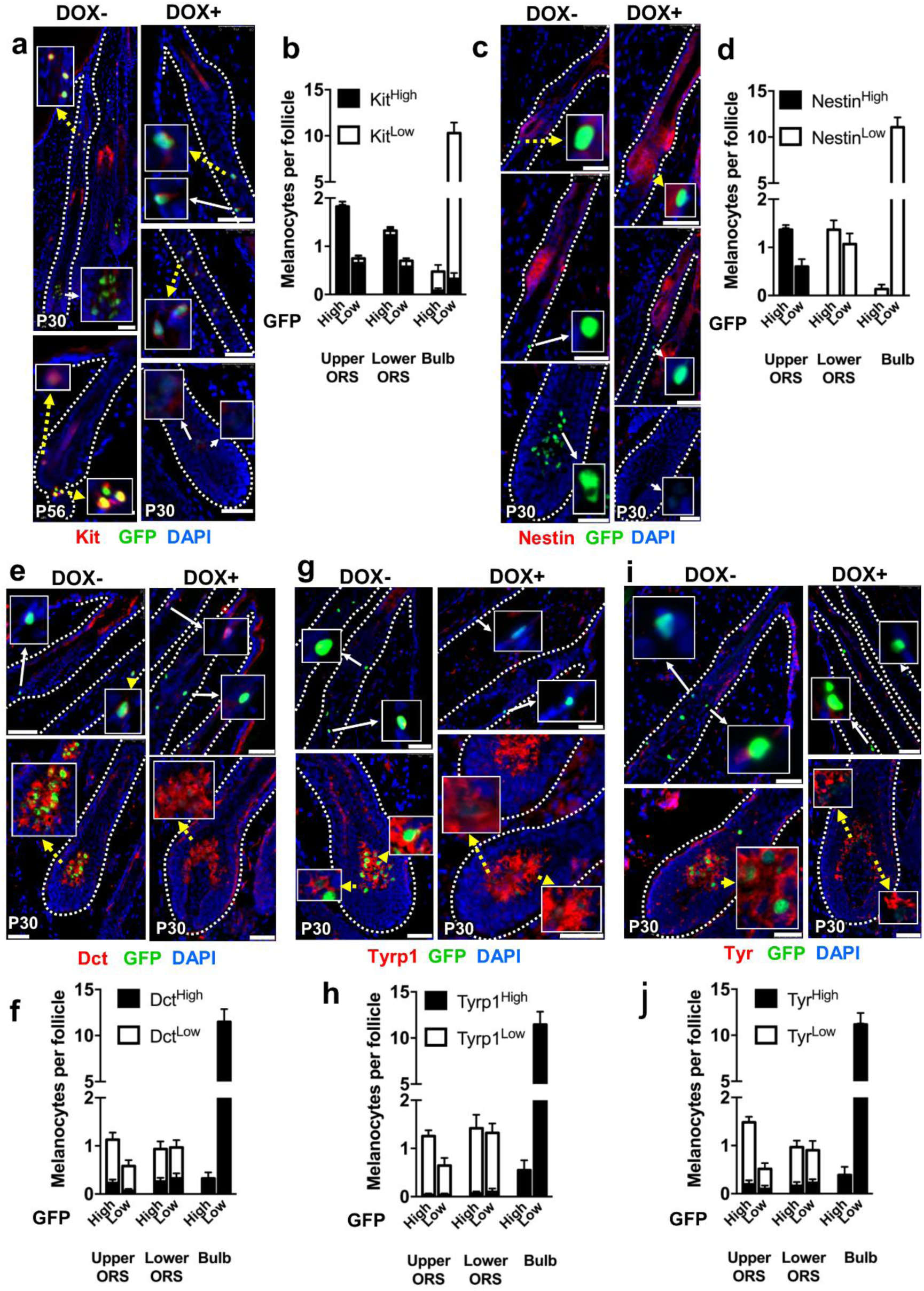
Kit and Nestin expression distinguish quiescent melanocytes while melanogenic markers characterize proliferating melanocytes in anagen HFs. Immunofluorescence and quantification of Kit (a, b), Nestin (c, d), Dct (e, f), Tyrp1 (g, h) and Tyr (i, j) expression in GFP^high^ and GFP^low^ melanocytes with (right panels) and without (left panels) DOX treatment. Top panels including middle panels in (a) and (c), show expression of the marker in upper and lower ORS. Bottom panels show expression in melanocytes of HF bulb at P30 (except (a) with image showing Kit expression at P56 telogen HF). Scale bars, 50µm. (mean ± SEM, minimum 30 HFs quantified/animal). Inset images identifies melanocytes with (yellow dotted arrow) and without (white arrow) expression of the concerned marker. Scale bars, 50µm. See also figures S6-S10 and table S4.

**Figure 5.**
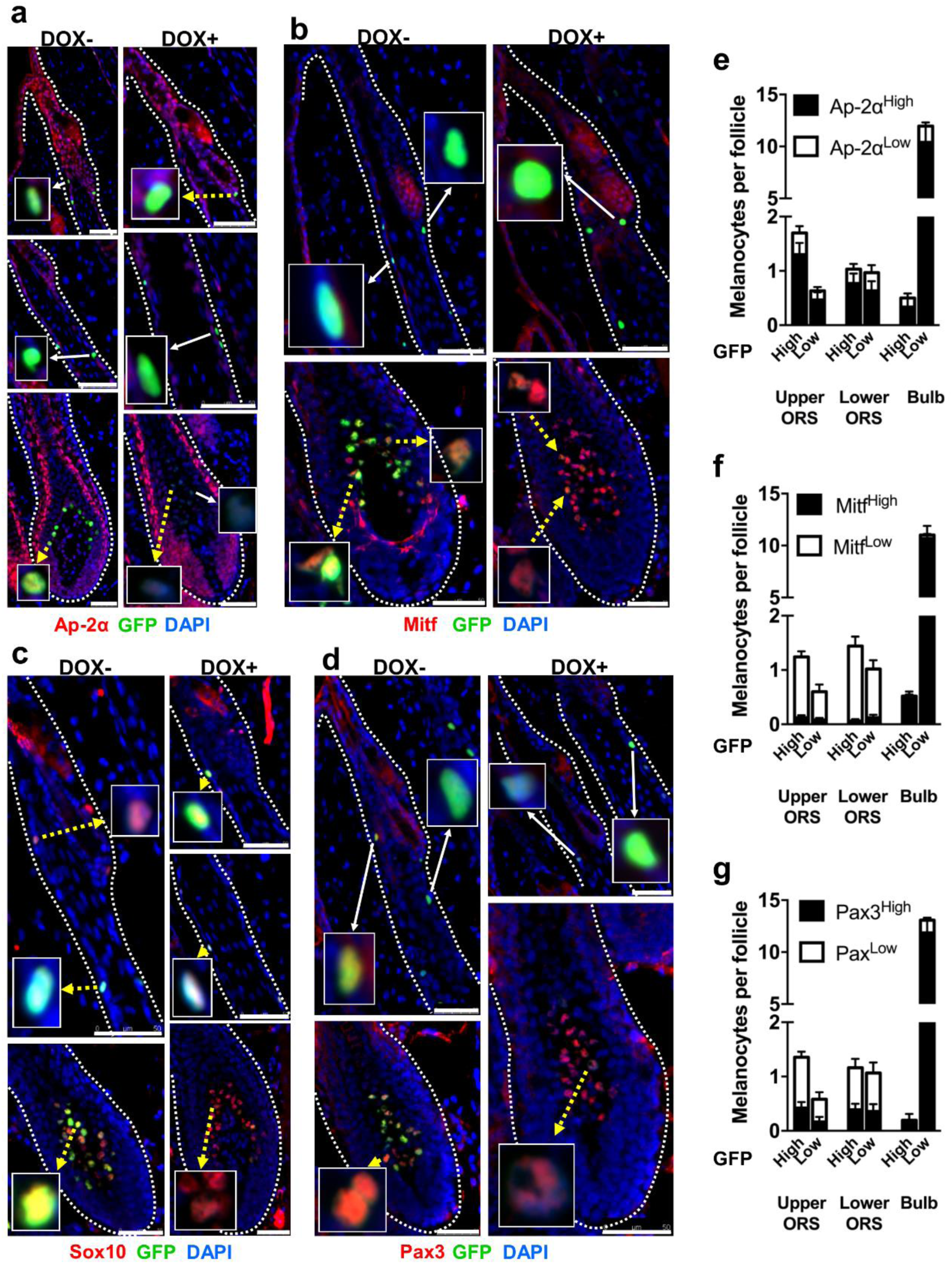
Transcription factors: Mitf and Pax3 is expressed only in proliferating melanocytes from HF bulb in late anagen. Expression of Ap-2α (a), Mitf (b), Sox10 (c) and Pax3 (d) in GFP^high^ and GFP^low^ melanocytes with (right panels) and without (left panels) DOX treatment during anagen onset. Top panels (and middle panels where applicable) shows expression in ORS region and bottom panels in bulb region. Inset images depict examples of melanocytes with (yellow dotted arrow) and without (white arrow) expression of the interested transcriptional factor. Quantification of Ap-2α (e), Mitf (f) and Pax3 (g) in GFP^high^ and GFP^low^ melanocytes from the images Scale bars, 50µm. (mean ± SEM, minimum 30 HFs counted/animal) See also figures S11- S14 and table S3c and S4.

### Ki67 expression and GFP retention in proliferating and quiescent melanocytes in anagen HFs

The heterogeneity of GFP expression with Dox in Sox10+ melanocytes suggested that anagen HF melanocytes proliferate differentially and dilute the GFP label. We hypothesized that GFP expression in GFP^low^ cells was diluted because they were more proliferative. To demonstrate this, we used another proliferation marker, Ki67, to correlate with GFP and its retention.

Ki67 expression was analyzed in P30 *Dct-*H2BGFP anagen HFs. In control mice that did not receive doxycycline, most Ki67+GFP+ cells resided in the HF bulb (Figure 2e), correlating strongly with the presence of GFP^low^ cells in Dox-administered mice (Figure 2b). There were fewer Ki67+ melanocytes in the HF ORS (Figure 2e) where we had previously observed a larger population of GFP^high^ cells (Figure 2c). Ki67 expression and localization was analyzed in HFs of P30 mice that received Dox from P19. Quantitatively, comparison of Ki67 and GFP^low^ distribution shows that 91% of GFP^low^ cells and 85% of Ki67+GFP+ cells are in the bulb (Table S3d) while <10% of Ki67+GFP^low^ melanocytes reside in the upper and lower ORS region of the anagen HF (Figure 2g). Similarly, almost all Ki67+GFP^high^ melanocytes were in the upper/lower ORS (6.2% and 4.1%, respectively, of 10.5% Ki67+GFP^high^ melanocytes; Figure 2g).

The correlation between Ki67 and GFP expression was further investigated by flow cytometry. The GFP^high^ cell population (Figure 3a) was significantly reduced from 0.43±0.05% in control mice to 0.21±0.10% in Dox-treated mice (Figure 3b,c; p-value=0.02), consistent with dilution of GFP expression following cell proliferation. Profound reduction in Ki67+GFP^high^ cells was observed (upper right, Figure 3d,e) from 0.23±0.08% to 0.03±0.01% after Dox treatment (Figure 3f;p=0.05). Ki67-GFP^high^ cells did not show a significant difference between samples from control (0.21±0.07%) and Dox-treated mice (0.19±0.02%;p-value=0.86), probably because of the larger size of this cellular subset. These results imply that the Ki67+GFP^high^ cell population in control mice that proliferated close to P30 shifted to the left due to GFP dilution. Selective loss of Ki67+GFP^high^ cells after Dox treatment is consistent with their property as a quiescent melanocyte population.

### Compartmental expression of McSC and melanocyte differentiation proteins during HF anagen

To further characterize anagen HF melanocytes in distinct HF compartments, we determined their relative expression of stem cell and differentiation proteins, including the McSC markers Kit and nestin. Kit, a receptor tyrosine kinase, has been characterized previously as a McSC marker (Harris et al. 2013; Nishimura et al. 2002) and is functionally important in melanocyte development (Mackenzie et al. 1997; Okura et al. 1995). Nestin identifies neuronal stem cells and other neural crest-derived pluripotent cells (Yaworsky and Kappen 1999), including select HF cells such as bulge/CD34+ McSCs reported previously (Amoh et al. 2005; Hoffman 2007; Joshi et al. 2019). Nestin is also expressed in keratin 15-negative (Krt15-) cells of the HF bulge. At P30, Kit expression was high in upper and lower ORS melanocytes, but not in bulb melanocytes. Within the ORS, the level of expression of Kit did not differ between GFP^high^ and GFP^low^ melanocytes (Figure 5a,b). In contrast, nestin expression was restricted to the upper ORS region, comparable to the bulge/LPP in telogen, and was invariantly high in GFP^high^ and GFP^low^ upper ORS cells.

We also determined the relative expression of melanocyte differentiation markers Dct, tyrosinase-related protein 1 (Tyrp1), and tyrosinase (Tyr). Analysis of their expression showed high levels of Dct, Tyr, and Tyrp1 in melanocytes in the HF bulb relative to ORS melanocytes (Figure 5e,g,i). There was no significant difference between their expression in quiescent GFP^high^ cells or proliferative GFP^low^ cells in any HF location (Figure 5f,h,j). Hence, the expression of melanocyte differentiation proteins is unrelated to the proliferative state of the melanocyte but is highly correlated with the presence of the cell in the anagen bulb.

### Compartmental expression of melanocyte transcription factors during anagen

Additionally, we analyzed expression of the transcription factors Ap-2α, Sox10, Pax3, and Mitf in quiescent melanocytes. Each has critical roles in melanocyte development and differentiation (Carreira et al. 2006; Carreira et al. 2005; Hornyak et al. 2001; Levy et al. 2006; Potterf et al. 2001; Seberg et al. 2017). Most P30 melanocytes expressed Ap-2α with no significant difference observed between GFP^high^ and GFP^low^ melanocytes in ORS and HF bulb (Figure 5a,e). Ap-2α was observed in non-melanocytic HF cells in the upper bulge and infundibulum (Figure 5a, upper), and in ORS cells approaching the bulb region (Figure 5a, lower), but was not observed in ORS cells between the bulge and the bulb (Figure 5a, middle panels). Mitf expression was most strongly correlated with differentiated melanocytes in the HF bulb, where its expression was invariant between GFP^high^ and GFP^low^ cells. In most upper ORS and lower ORS cells, Mitf was expressed at a low level (Figure 5b,f). Pax3 expression was similar to Mitf, with high levels of expression detected in GFP^high^ and GFP^low^ melanocytes in the bulb and lower fractions of cells in the upper/lower ORS expressing high Pax3 (Figure 5d,f). Sox10 expression was highly specific for melanocytes in the HF. Quantification of its relative expression showed no difference between quiescent and proliferating melanocytes (Figure 5c), consistent with the prior report that its expression can be detected at all levels of the HF during anagen (Harris et al. 2013).

## DISCUSSION

Our results demonstrate that anagen ORS melanocytes can be divided into Nstn^high^Kit^high^Mitf^low^Pax3^low^ and Nstn^low^Kit^high^Mitf^low^Pax3^low^ subsets corresponding to cells in the bulge/LPP and the lower ORS, respectively. Melanocytes in the anagen bulb instead have a Kit^low^Mitf^high^Pax3^high^ phenotype. None of these markers distinguishes between quiescent and proliferative ORS cells during anagen.

These results establish definitively the presence of quiescent melanocytes in anagen outside of the bulge/LPP region. In previous work using a depilation-induced anagen model (Botchkareva et al. 2001), only a fraction of ORS and bulb melanocytes expressed Ki67 in early- and mid-anagen. However, no label-retaining technique was used to confirm that Ki67-cells remained quiescent throughout anagen. Telogen bulge melanocytes expressed Dct alone and were Ki67-, consistent with our findings, and ORS melanocytes expressing Dct and a low level of Tyrp1 differed from fully differentiated melanocytes in the HF matrix expressing Dct/Tyrp1/Tyr. Their description of Kit expression differed from ours, whereby Kit expression in the bulge was variable during telogen, decreased or extinguished in this region during depilation-induced anagen, but high in the anagen matrix. Our findings are more consistent with a more recent study (Harris et al. 2013) which found that Kit is maintained in bulge/LPP and ORS cells in mid-anagen, with the percentage of Kit^high^ cells declining only slightly in telogen. The prior study used an antibody to Kit requiring tyramide amplification, possibly explaining the difference between their conclusions and those shared by us and the more recent study (Harris et al. 2013).

Although the bulge/LPP remains intact throughout the HF cycle, the SHG is transient and disappears as a recognizable structure following anagen onset. Hence the regeneration of McSCs in the SHG after the first anagen is dependent upon their appearance from another structure. Since neither bulge nor McSCs proliferate during telogen (Figure 1b,c), it is unlikely that bulge McSCs divide to repopulate the SHG during this stage, although our results do not formally exclude proliferation-independent interchange of McSCs between compartments. Instead, we propose that quiescent melanocytes outside the bulge/LPP during anagen may represent a source for SHG regeneration during telogen, with the survival of select quiescent cells throughout anagen and catagen reforming SHG McSCs. Identifying markers which distinguish quiescent anagen melanocytes will be important to conduct fate-mapping and other experiments required to test this hypothesis directly.

## MATERIALS AND METHODS

### *In Vivo* labeling of melanocytes

*Dct*-H2BGFP mice (Joshi et al. 2018) were used to identify melanocytic cells. To label proliferating cells, *Dct*-H2BGFP mice were intraperitoneally injected with 10mM BrdU (50µg/g body weight, Molecular Probes) every 12h from P21-P30 or for one-week periods during telogen. Control littermates were injected with PBS. Skin samples were collected from 3 different regions of dorsal skin-frontal, lateral and hind at P30, P56, P63 and P70 following BrdU administration during anagen or telogen. *Dct*-H2BGFP mice were administered doxycycline hyclate (Sigma; 2g/l dissolved in drinking water; changed every 48h) from P19 to P30 to identify GFP-retaining cells. The control group received regular drinking water. Dorsal skin samples were collected during late anagen at P30 (n=3). All experiments were conducted under the auspices of an approved animal care protocol of the University of Maryland School of Medicine.

### Immunofluorescence analysis

10µm cryosections were cut from OCT (Tissue-Tek)-embedded skin and fixed with 4% paraformaldehyde. For BrdU labeling identification, sections were fixed with 4N HCl (30m) and 0.1M sodium borate wash. The sections were permeabilized and blocked (10% FBS/1% BSA/0.1% Triton-X) for 1h. MOM kit (Vector laboratories) was used for mouse primary antibodies, with overnight incubation followed by secondary antibody incubation (1h). Section mounting solution contained DAPI (Vector laboratories).

Immunofluorescence images were obtained by Olympus upright or DMi8 Leica microscopes with imaging software (Slidebook/Leica Application Suite X). For BrdU detection, images were taken within 24h of fixation to prevent fluorescence loss. Raw counting data is in tables S1-6.

### Cell sorting analysis

Skin from P30 *Dct*-H2BGFP mice with/without Dox treatment was trypsinized in 0.5% trypsin (Affymetrix) at 37°C (15min) after removal of subcutaneous fat, cut into small fragments, and incubated in thermolysin-low Liberase (Roche) at 37°C (1h) followed by treatment with 0.05% DNase (Sigma) and 5% FBS. Fragments were dissociated into single cells by repeated passage through a 60ml syringe. Remaining impurities and clumps were filtered out with 100µm cell strainers (Fisher) and 40 µm cell strainers (Fisher). The final cell suspension was prepared in 5% FBS.

Dead cells in the cell suspension were identified by adding 7-AAD (BD Biosciences). For Ki67 flow cytometry, fixable viability dye eFluor 780 (30min) was used to label non-viable cells followed by fixation with 4% paraformaldehyde solution (10min) and blocking in 10% FBS/1% BSA/ 0.1% Triton-X (30m). Cells were incubated in AlexaFluor 647 conjugated Ki67 antibody (1:20, rabbit mAb, clone D3B5, Cell Signaling). Flow cytometry data was acquired by BD-LSRII flow cytometer.

### Statistical analysis

Two-way unpaired Student’s t-test (Figure 1c) or one-way ANOVA (Figure 1e,2d) was used to compare BrdU+ and BrdU– cells as well as GFP^high^ and low GFP^low^ cells (Prism). Pearson’s chi-squared test was used to confirm correlation between Ki67 expression and GFP^low^ cells after Dox treatment (Figure 2g). The flow cytometry data was analyzed by Flowjo software followed by two-way unpaired Student’s t-test (Figure 3c,f).

