## Supplementary Figures and Tables for "Complementary *In Vivo* Approaches Reveal Quiescent Melanocytes in Anagen Outside the Hair Follicle Bulge"

**Supplementary Figure 1.** (a) Characterization of HF stages by alkaline phosphatase (AP) staining. The left image shows AP stained dermal papillae enclosed by ORS characteristic of anagen HF in the skin cross section of our BrdU administered mice. The P49 mice from the litter used for telogen BrdU labeling show exposed dermal papillae suggesting onset of second telogen stage. (b) BrdU incorporation in melanocytes of second telogen to anagen transition HFs at P70. Appearance of BrdU+ melanocytes was observed in follicles transitioning in anagen. (c) Quantification of BrdU incorporation in the anagen HFs at P70. The highest number of BrdU+ GFP+ cells were present at the bulb (mean ± SEM, n = 3, control = 3).

**Supplementary Figure 1**

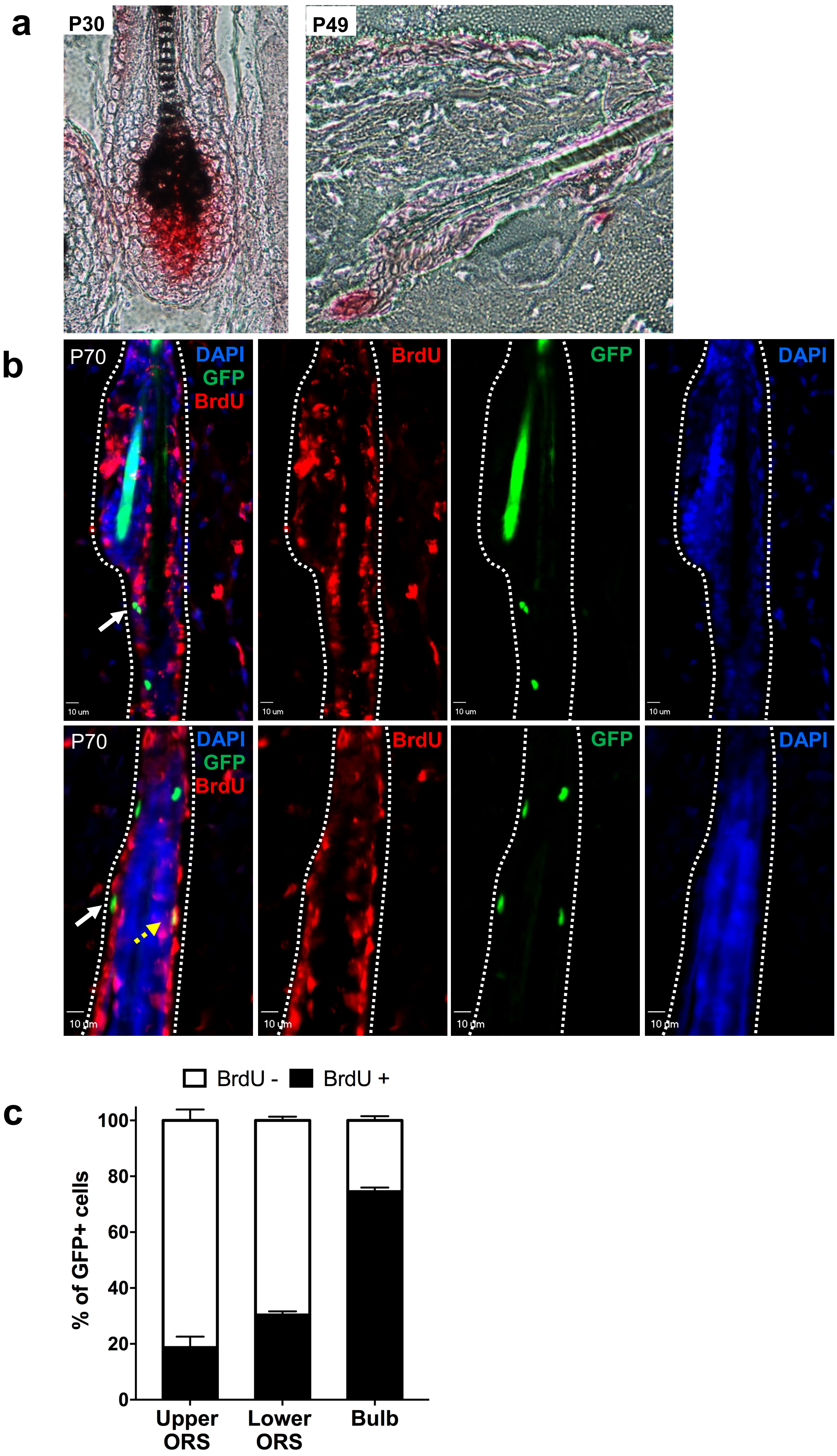

**Supplementary Figure 2.** BrdU labeling in murine HFs. (a) Immunofluorescence images of Rare BrdU- GFP+ Melanocyte observed near sebaceous gland above bulge region in telogen HF. (b) Immunofluorescence images of McSCs at bulge and SHG of earlier, mid and late telogen using an alternative mouse monoclonal antibody (clone B44, BD bioscience). Quantification of BrdU labeling show that 99.2±0.5% of SHG McSCs and 100% of bulge McSCs from P49 to P56, 98.8±0.1% of SHG McSCs and 96.1±1.8% of bulge McSCs from P56 to P63, and 97.7±0.8% of SHG McSCs and 97.6±0.2% of bulge McSCs were BrdU-. (mean ± SEM, n = 3, control = 3) (c) Immunofluorescence of BrdU labeling using the alternative mouse monoclonal anti-BrdU antibody shows BrdU– melanocytes in both upper and lower ORS of anagen HF (mean ± SEM, n = 3, control = 3).

**Supplementary Figure 2**

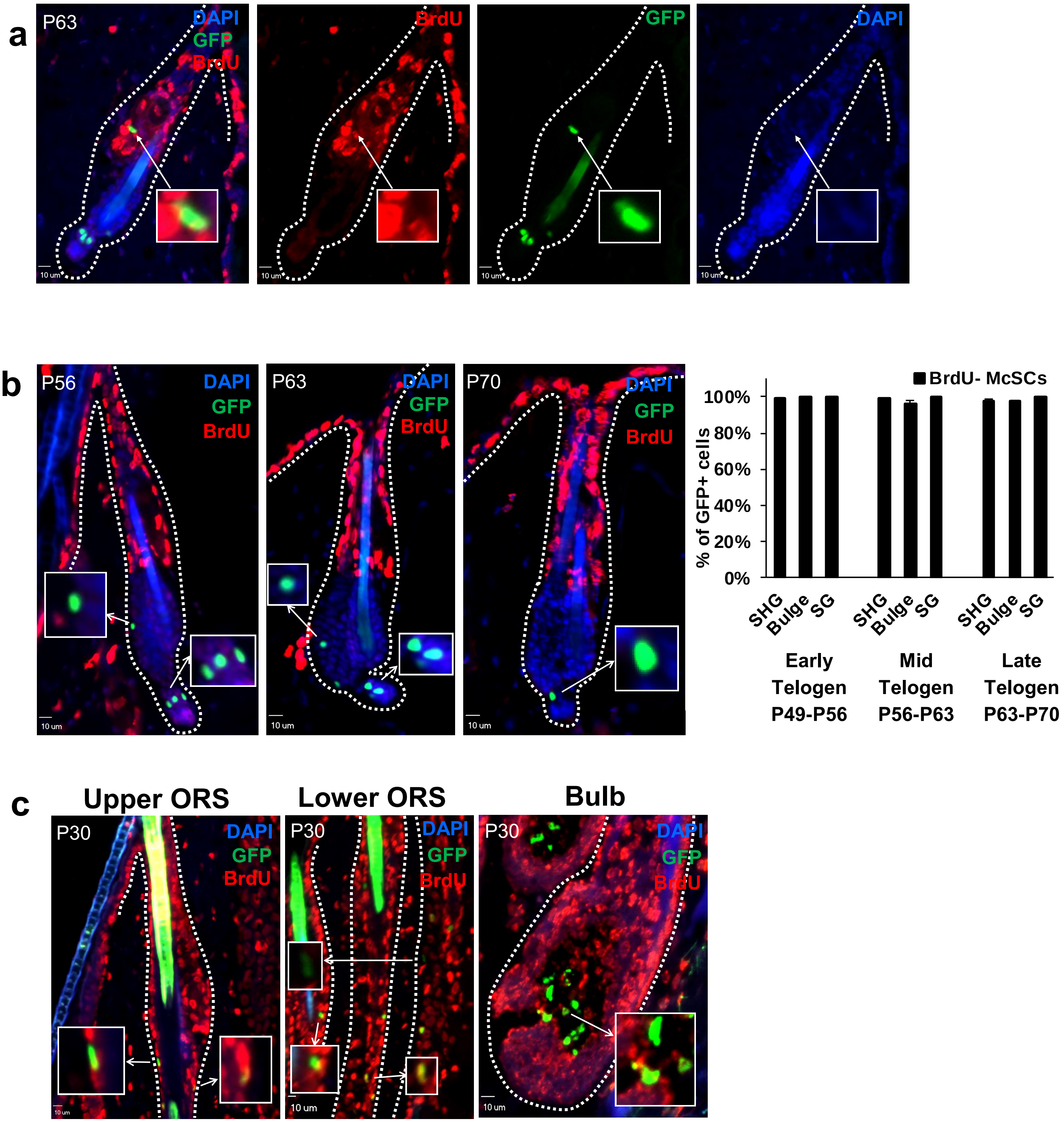

**Supplementary Figure 3.** Distinction between $GFP^{high}$ and $GFP^{low}$ melanocytes in anagen HFs at P30 after doxycycline treatment. (a) The figure shows examples of $GFP^{high}$ and $GFP^{low}$ melanocytes with their calculated corrected total cell fluorescence (CTCF). Melanocytes with CTCF value of 4000 were considered $GFP^{low}$. (b) Quantification of immunofluorescence images represented in Figure 2b and 2c for total GFP+ melanocytes classified based on its location in upper ORS, lower ORS and bulb. Graph shows significantly higher number of GFP+ cells in HFs from control mice (mean ± SEM, n = 3, control =3) $^{**}P < 0.01$ between GFP+ melanocytes in doxycycline treated vs control mice (Student's t-test). (c) Quantification of immunofluorescence images shown in Figure 6c for SOX+ GFP+ and SOX10+ GFP- cells categorized based on location in upper ORS, lower ORS and bulb. Graph shows significantly higher number of SOX+GFP- cells in HFs from doxycycline treated mice compared to control mice. (mean ± SEM, 24 HFs counted) $^{****}P < 0.0001$ between doxycycline treated vs control mice for SOX+GFP- cells (Student's t-test).

Supplementary Figure 3

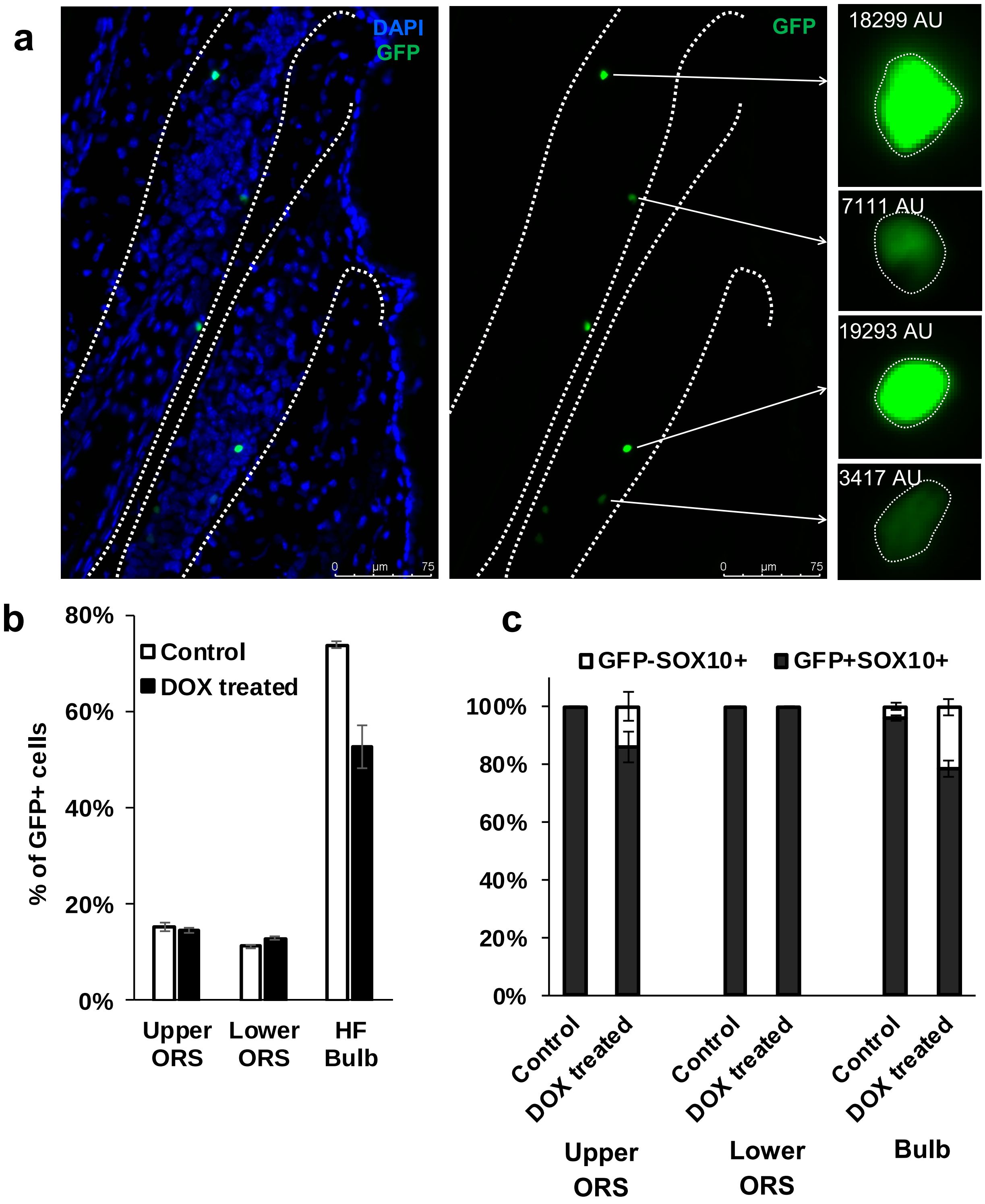

**Supplementary Figure 4.** Immunofluorescence images of CD34 expression in melanocytes during anagen. CD34 expression was observed only in bulge area of the anagen HF with or without DOX treatment (first and third row). Melanocytes in this region colocalized with CD34 (first row, left inset image; third row, left inset image). Cd34 expression was absent in lower ORS (third row, right inset image) or in the HF bulb (second and fourth row).

**Supplementary Figure 4**

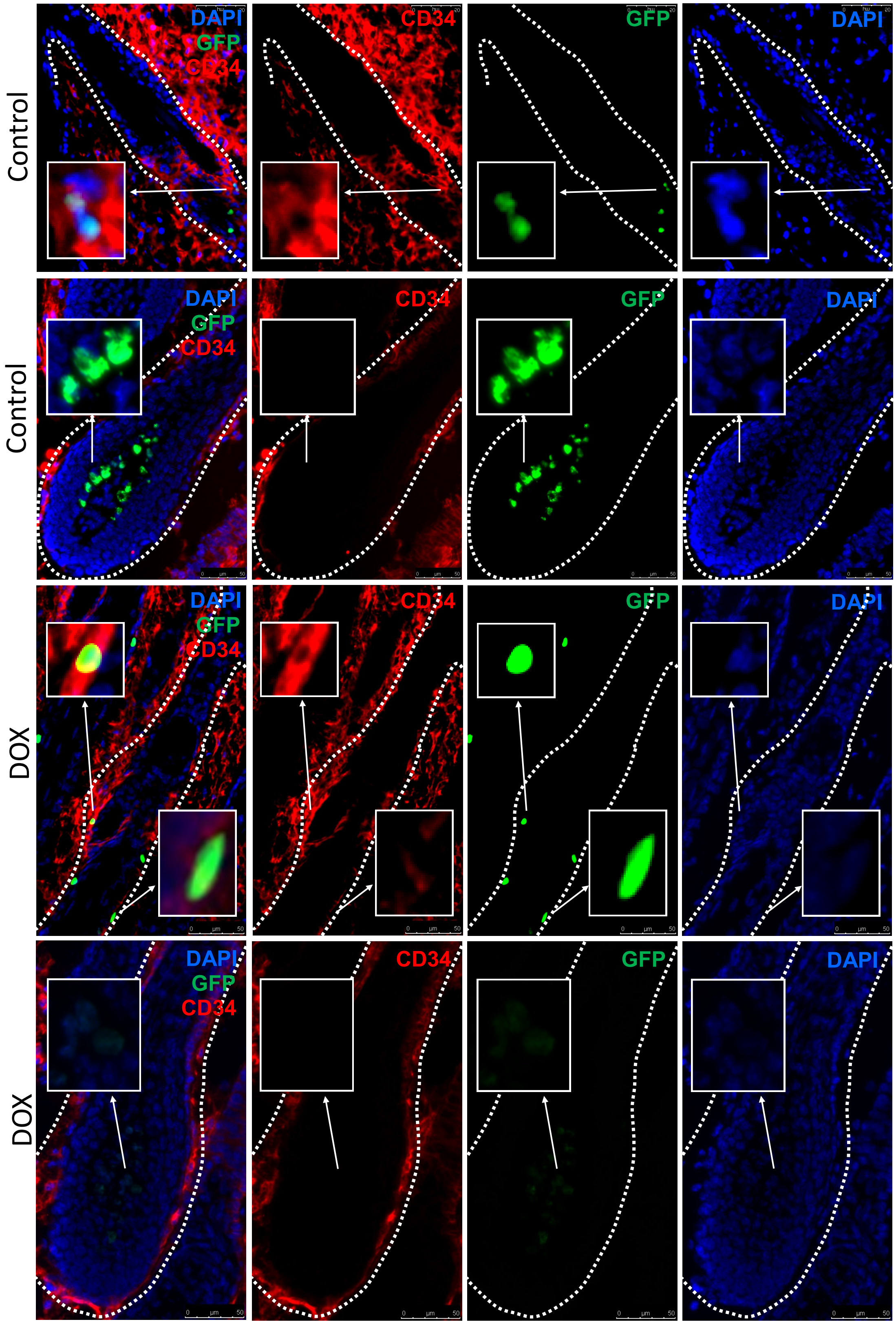

**Supplementary Figure 5.** Single channel immunofluorescence images of Figure 1d showing BrdU labeling in GFP+ melanocytes of P30 HFs from mice administered BrdU from P21 to P30.

Supplementary Figure 5

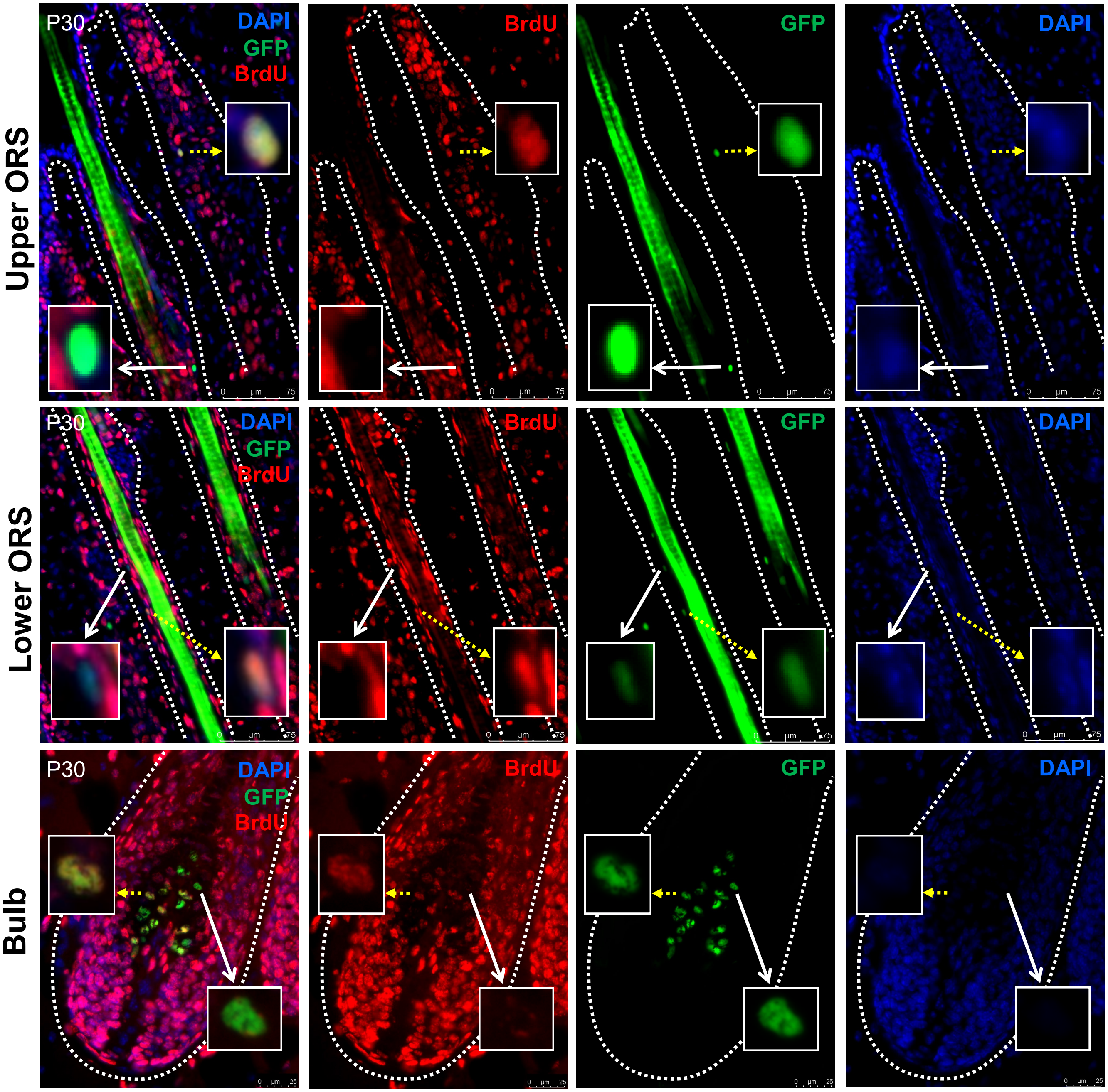

**Supplementary Figure 6.** Single channel immunofluorescence images of Figure 1b showing BrdU labeling of HFs during early, mid and late telogen.

**Supplementary Figure 6**

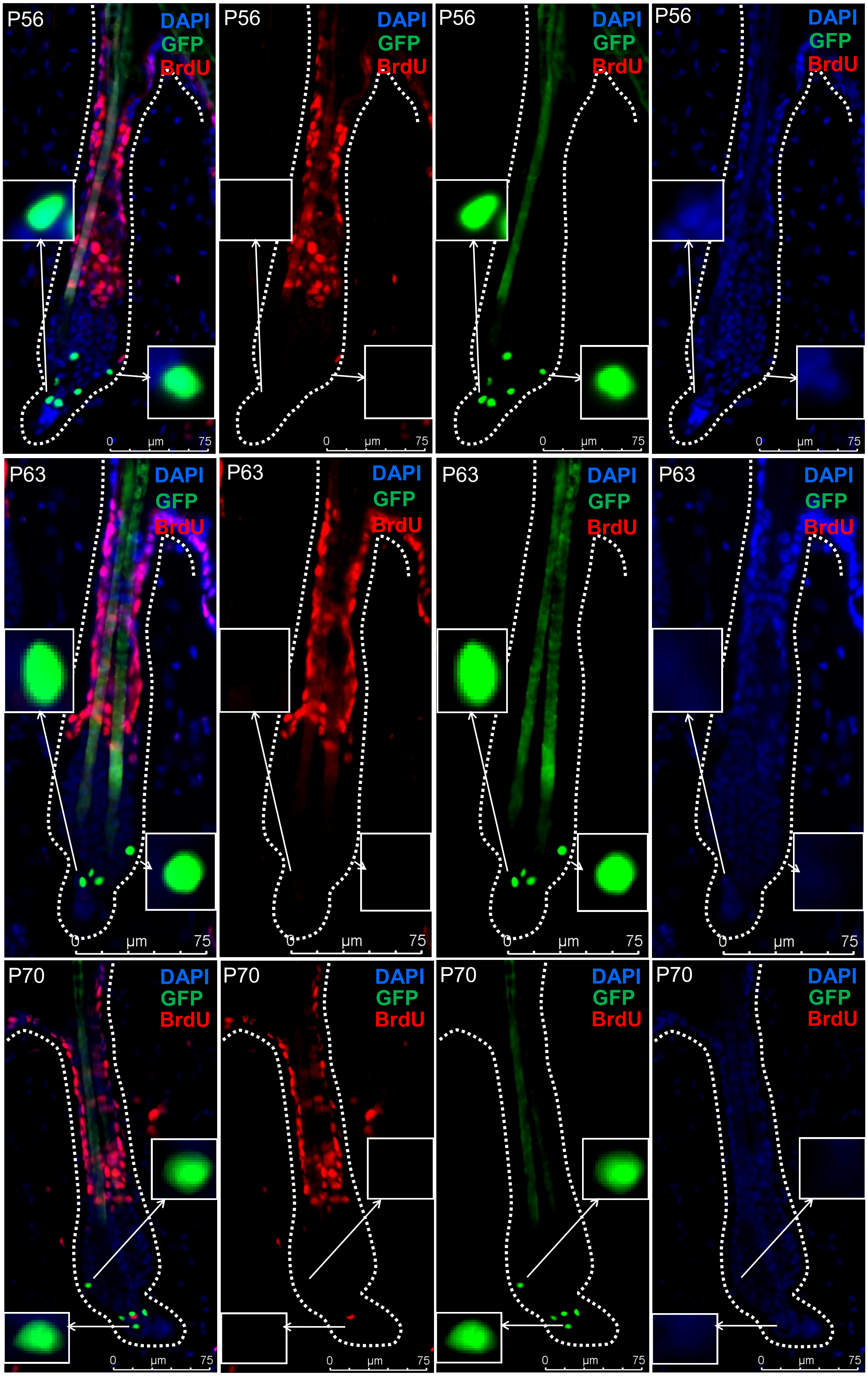

**Supplementary Figure 7.** Single channel Immunofluorescence images of Figure 4a (Kit expression +/- DOX treatment during anagen).

**Supplementary Figure 7**

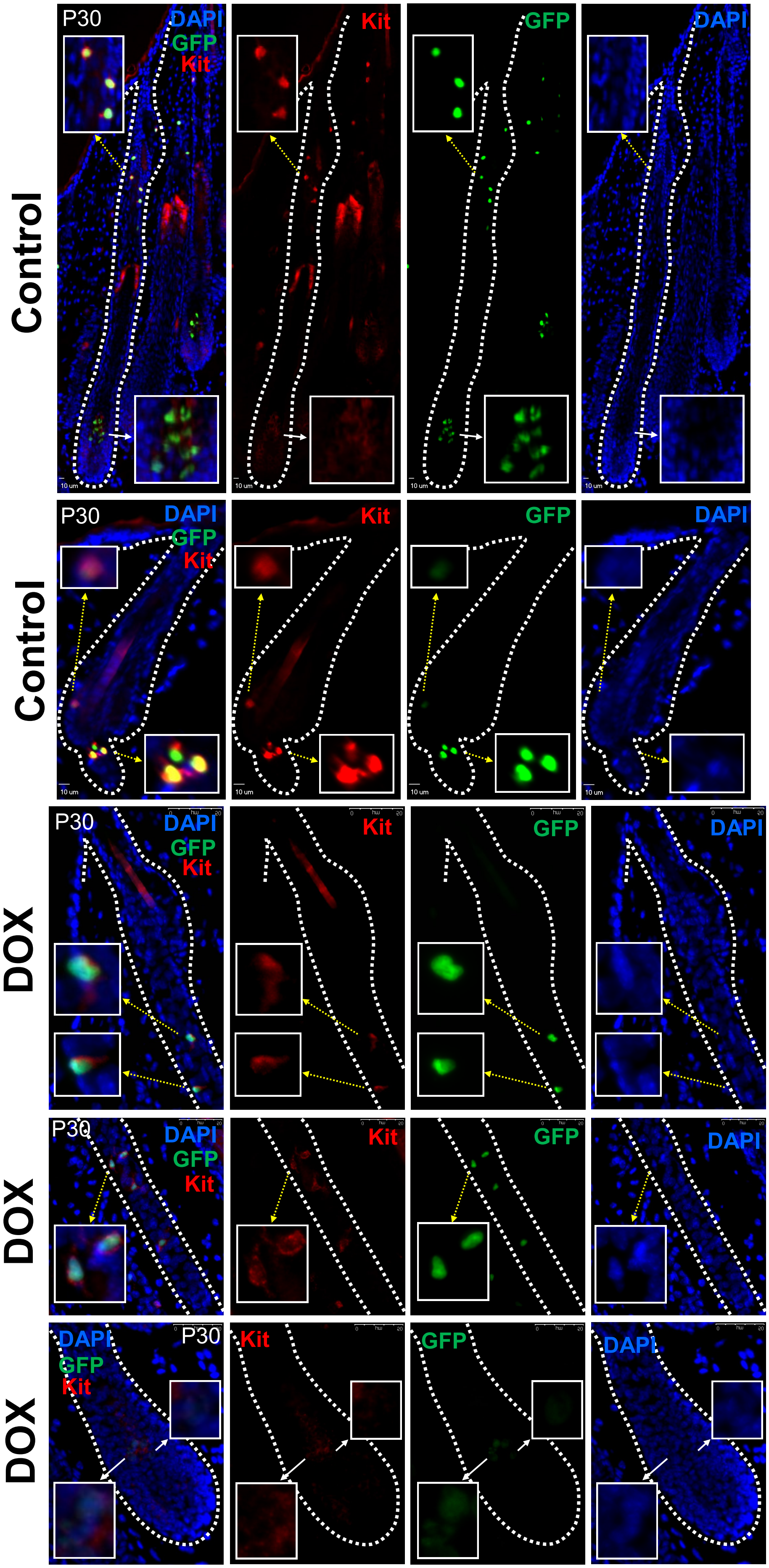

**Supplementary Figure 8.** Single channel immunofluorescence images of Figure 4c (Nestin expression +/- DOX treatment during anagen).

**Supplementary Figure 8**

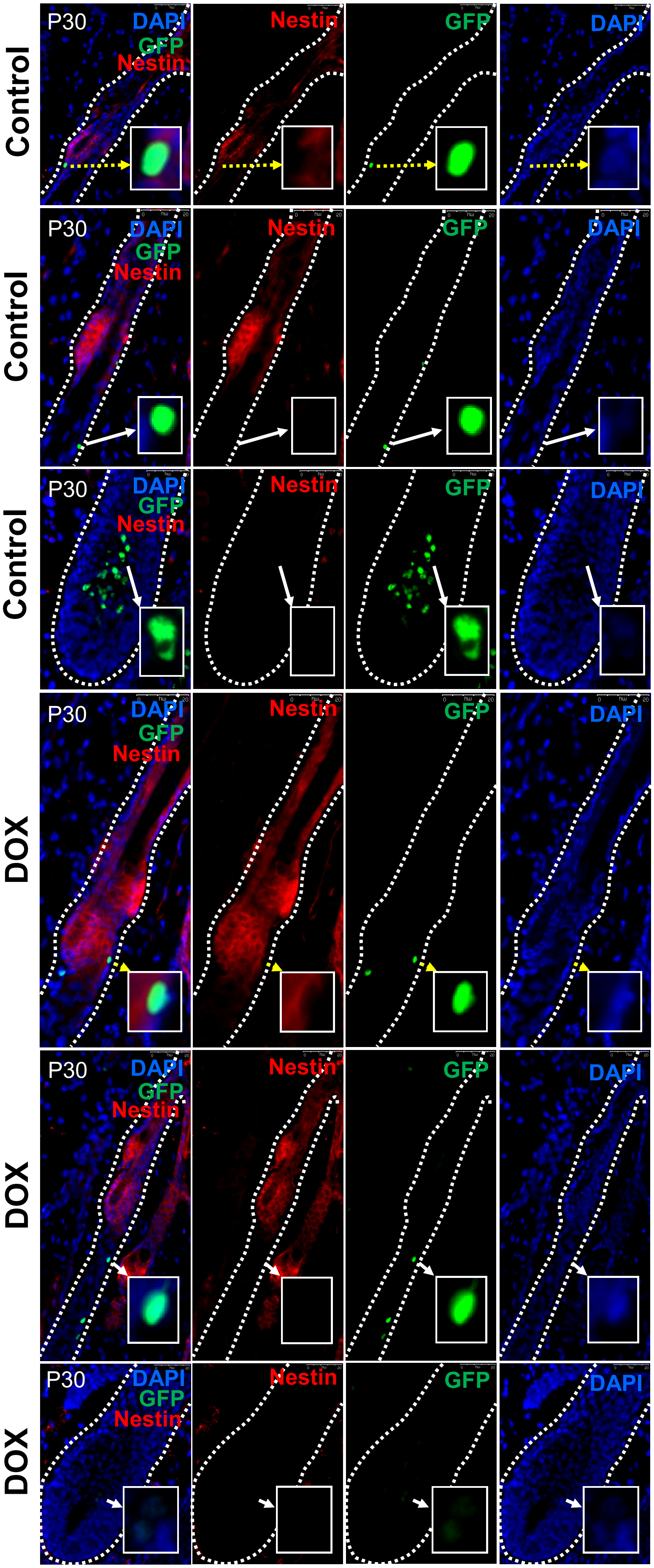

**Supplementary Figure 9.** Single channel immunofluorescence images of Figure 4e (Dct expression +/- DOX treatment during anagen).

**Supplementary Figure 9**

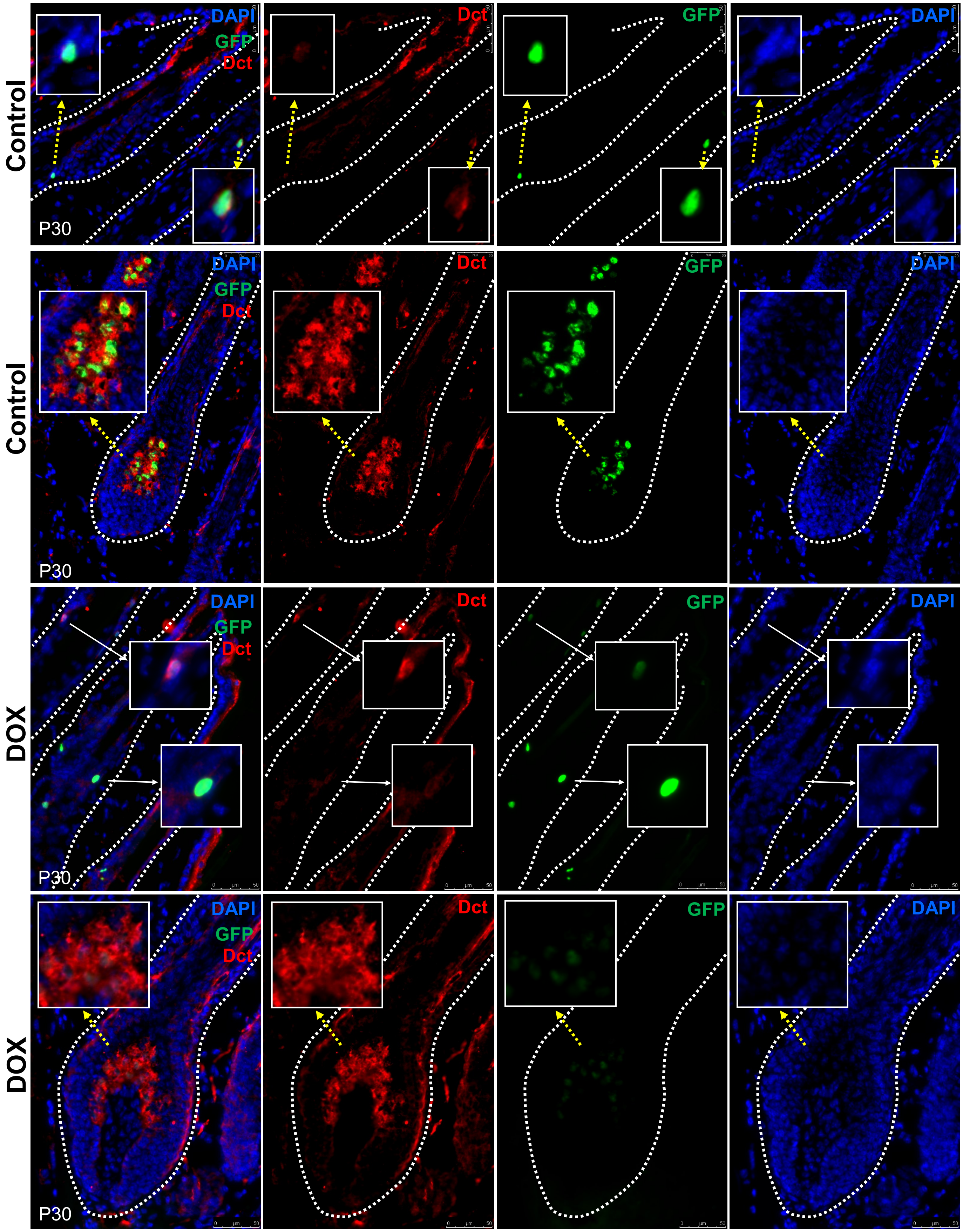

**Supplementary Figure 10.** Single channel immunofluorescence images of Figure 4g (Tyrp1 expression +/- DOX treatment during anagen).

**Supplementary Figure 10**

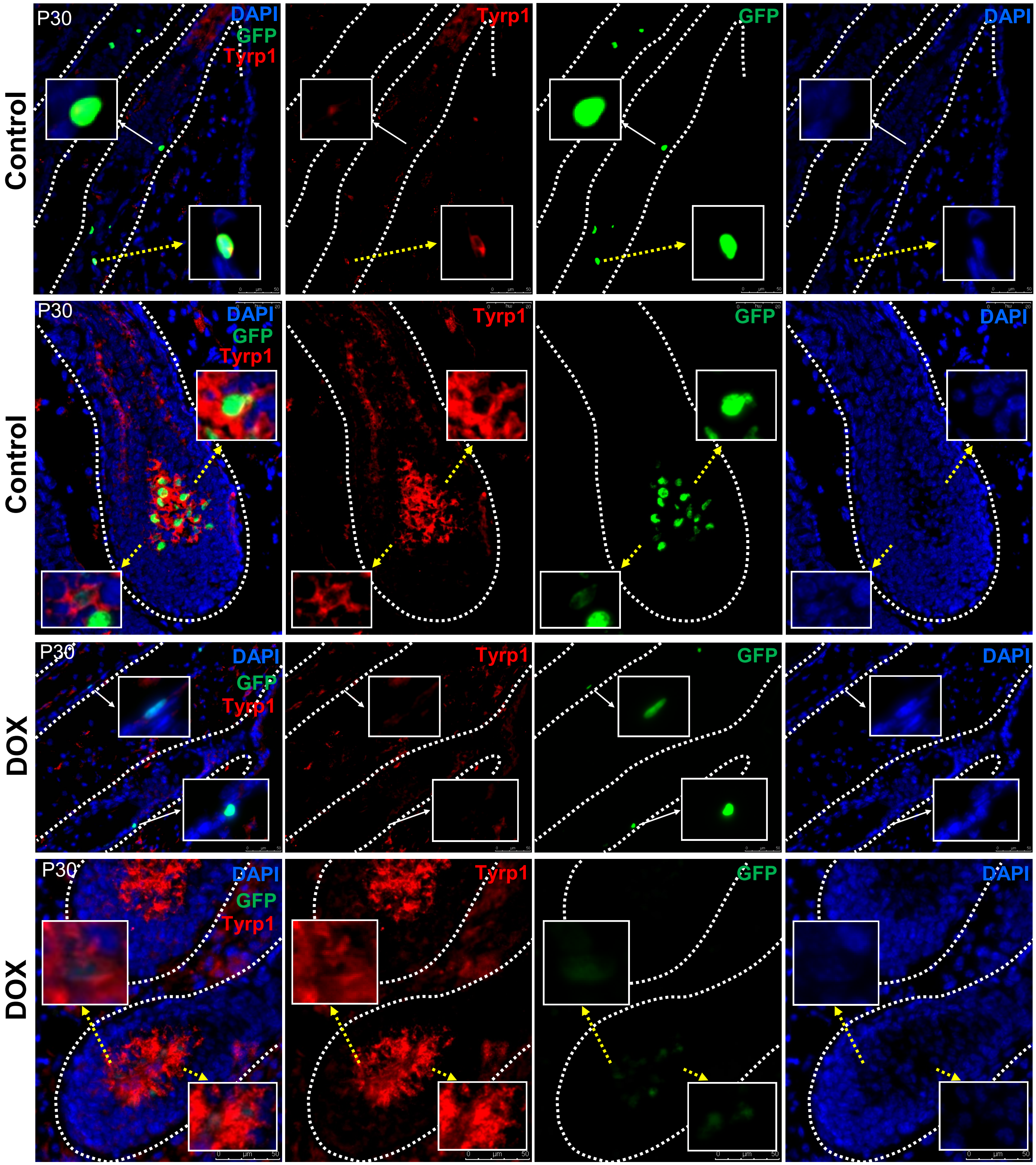

**Supplementary Figure 11.** Single channel immunofluorescence images of Figure 4i (Tyr expression +/- DOX treatment during anagen).

**Supplementary Figure 11**

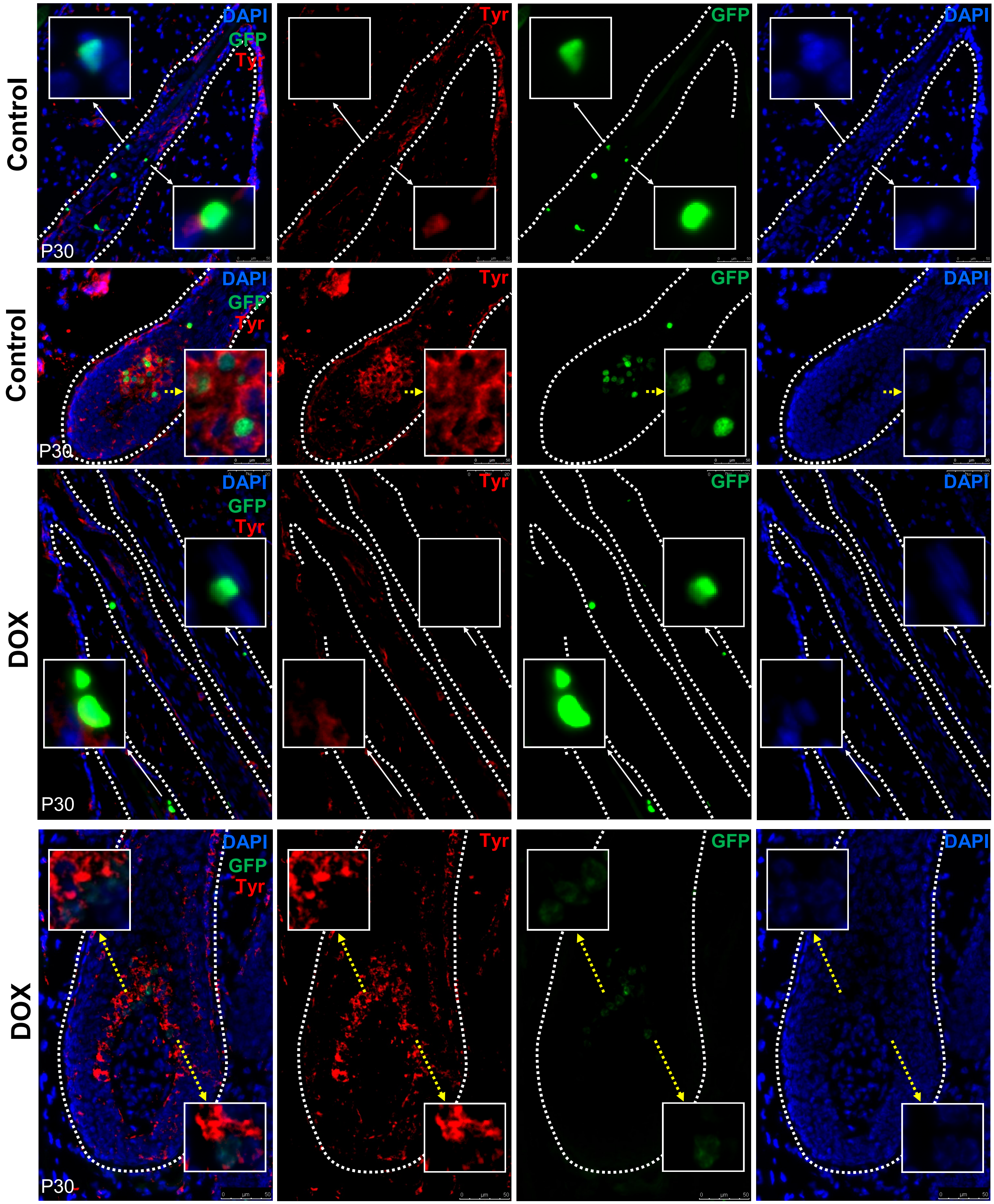

**Supplementary Figure 12.** Single channel immunofluorescence images of Figure 5a (Ap-2α expression +/- DOX treatment during anagen).

Supplementary Figure 12

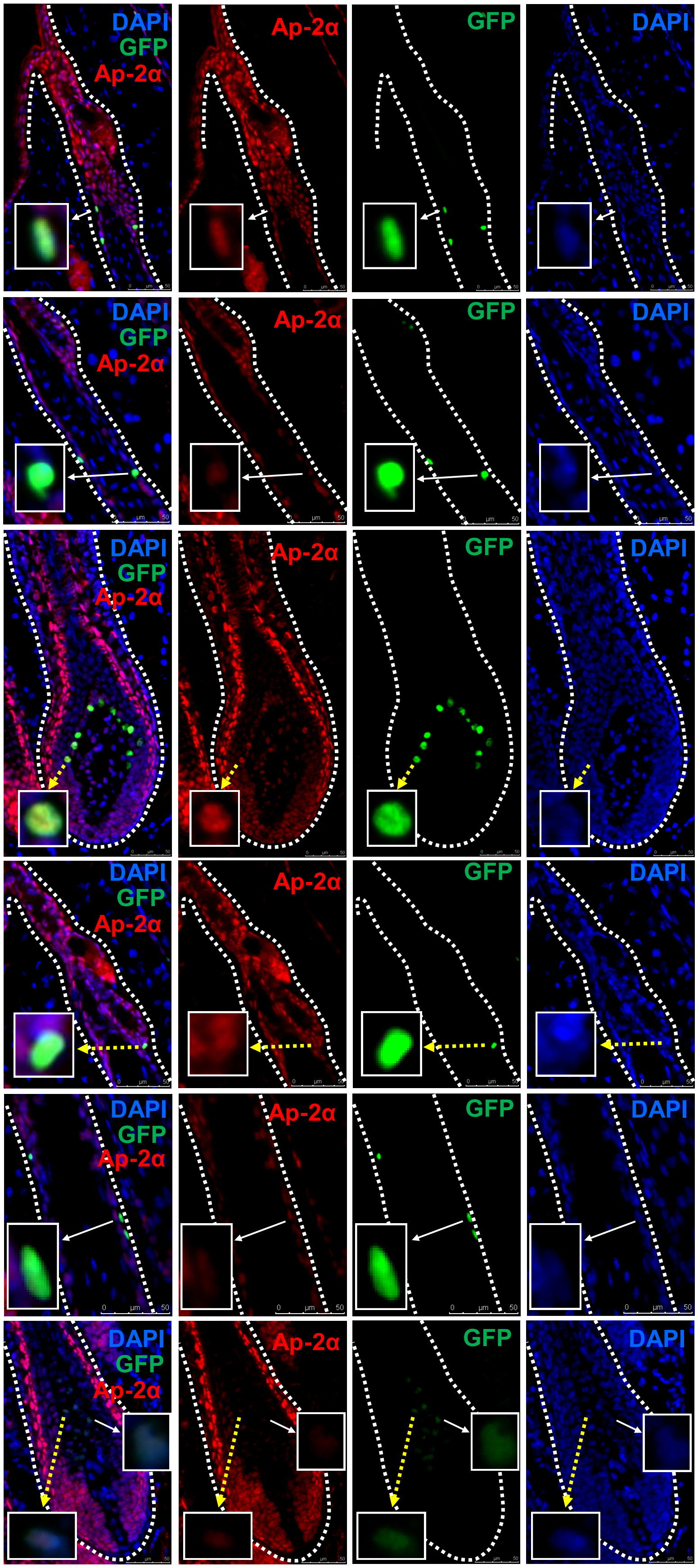

**Supplementary Figure 13.** Single channel immunofluorescence images of Figure 5c (Sox10 expression +/- DOX treatment during anagen).

**Supplementary Figure 13**

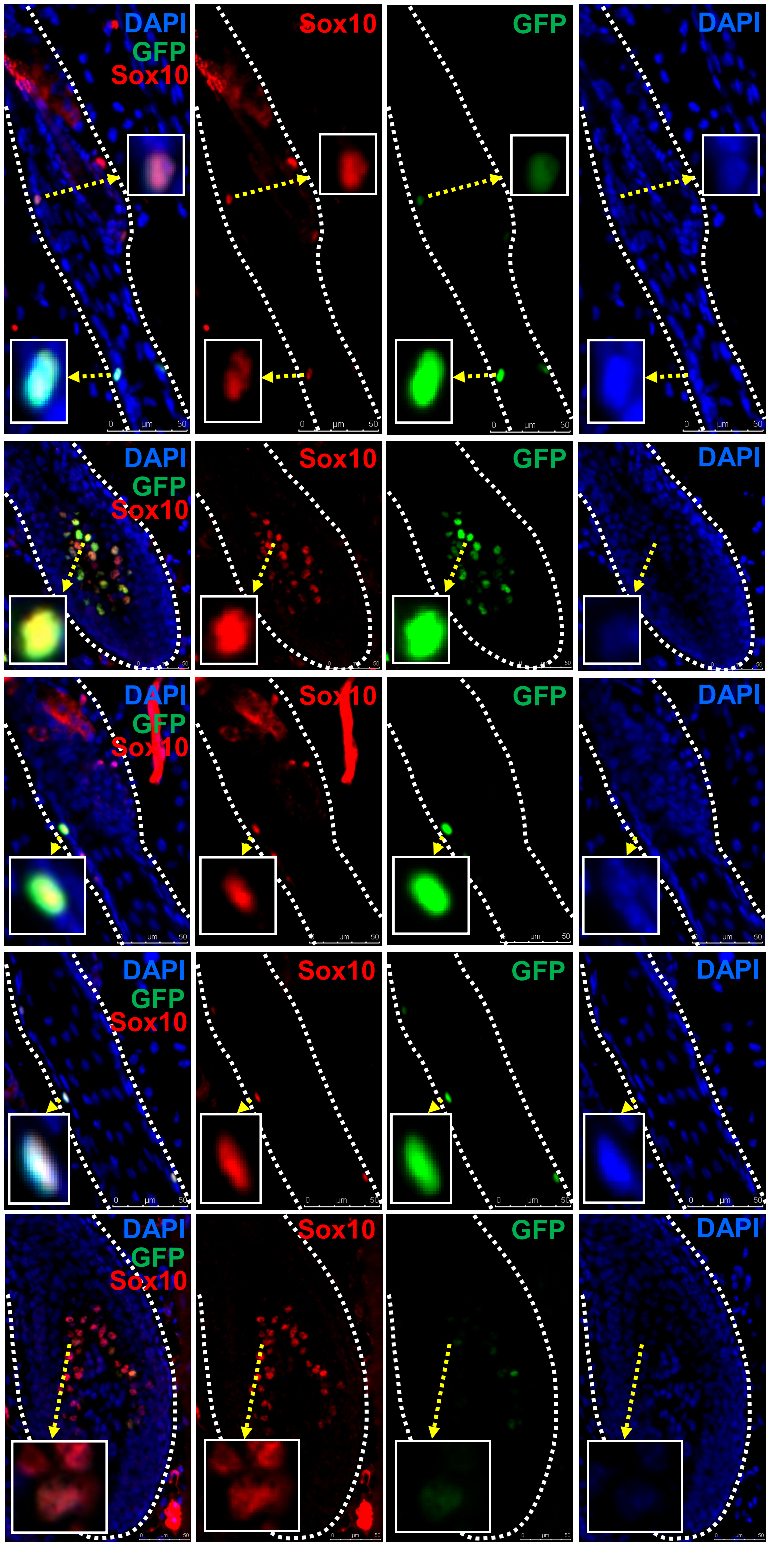

**Supplementary Figure 14.** Single channel immunofluorescence images of Figure 5d (Pax3 expression +/- DOX treatment during anagen).

**Supplementary Figure 14**

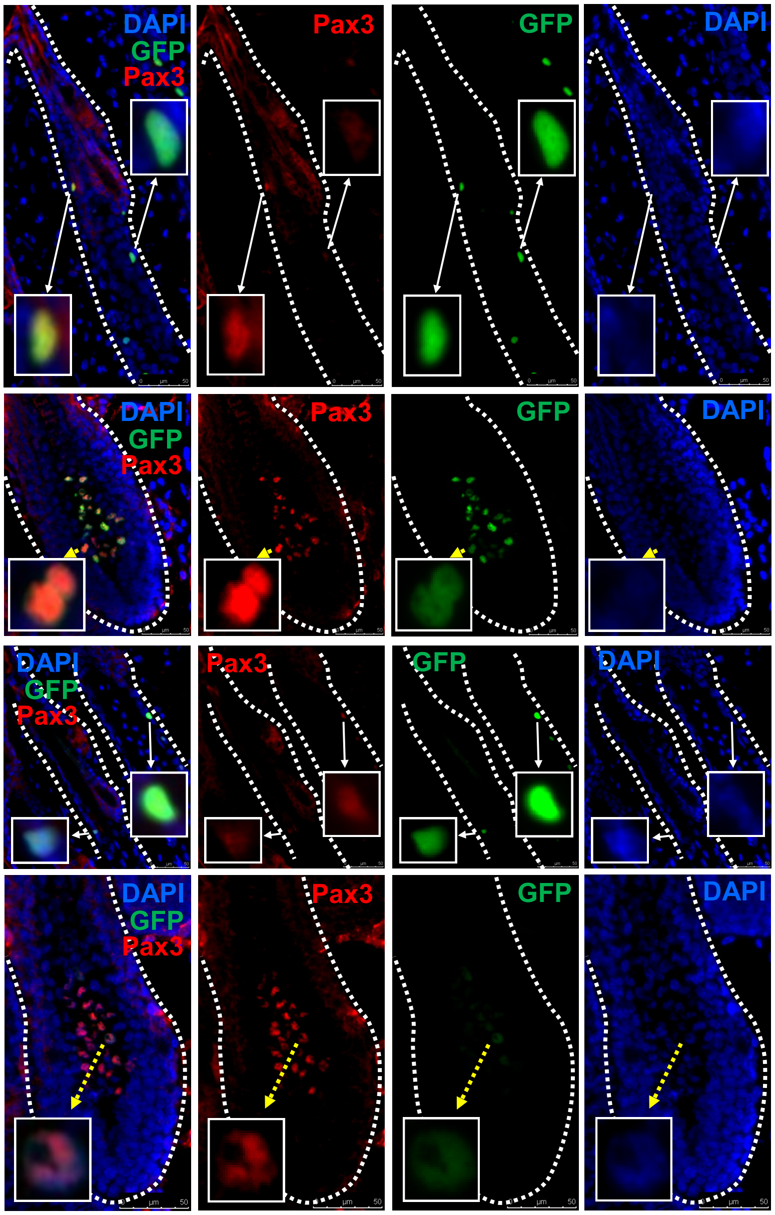

**Supplementary Figure 15.** Single channel immunofluorescence images of Figure 5b (Mitf expression +/- DOX treatment during anagen).

**Supplementary Figure 15**

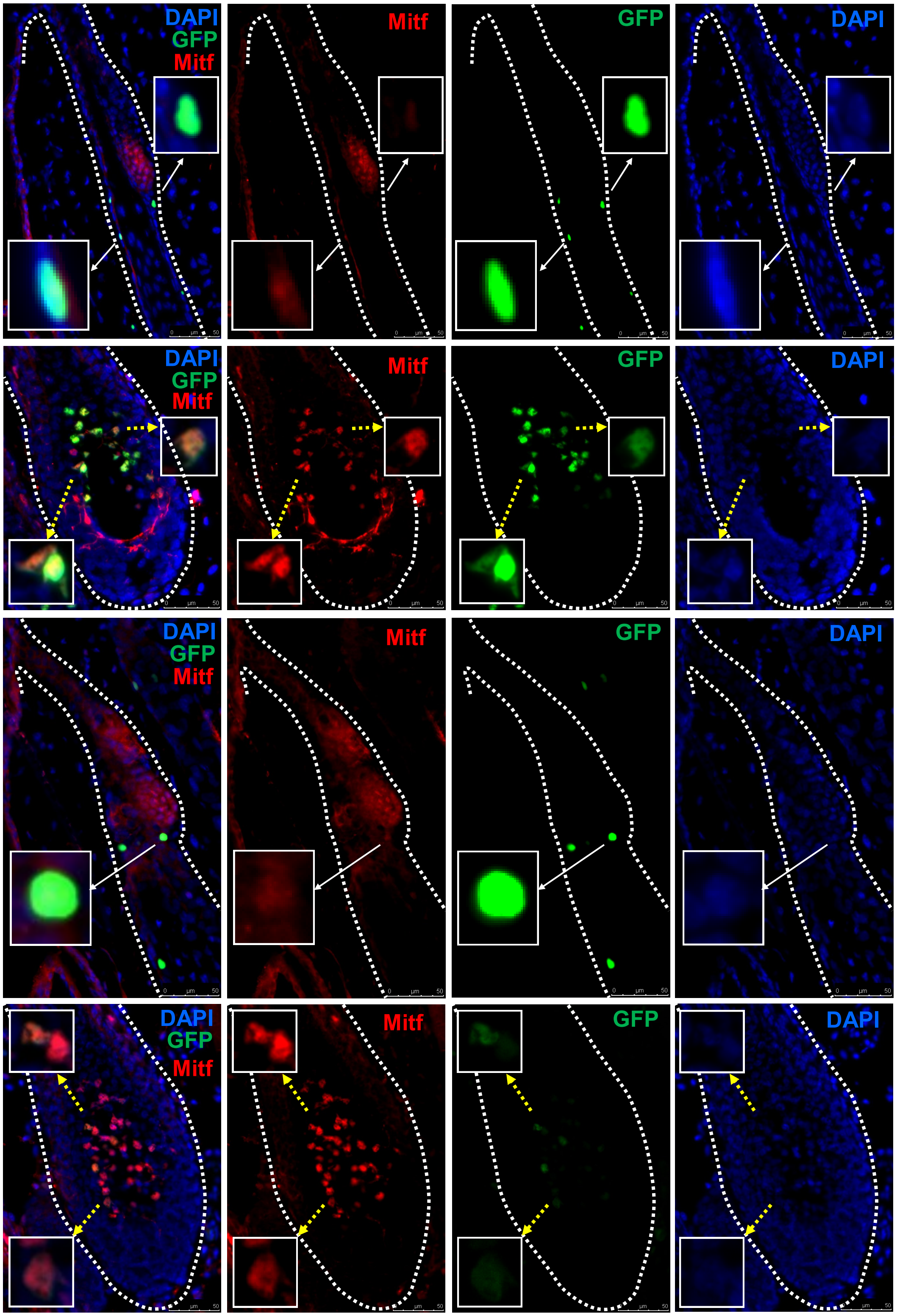

**Supplementary Table 1.** Quantification of BrdU labeling during telogen (P49-P56, P56-P63, P63-P70) (top table), the third anagen where some anagen follicles were observed at P70 (middle table) or second anagen (P21-P30). The table lists the total number of HFs counted and total BrdU+ and BrdU - cells from each mouse. (n=3)

**Supplementary Table 1**

**Second Telogen**

| Treatment period | Sample | No. of follicles | SHG | | Bulge | | SG | |
|---|---|---|---|---|---|---|---|---|
| | | | BrdU+ | BrdU- | BrdU+ | BrdU- | BrdU+ | BrdU- |
| P49-P56 | BrdU treated mouse 1 | 100 | 0 | 232 | 0 | 76 | 0 | 9 |
| | BrdU treated mouse 2 | 100 | 11 | 235 | 4 | 88 | 0 | 12 |
| | BrdU treated mouse 3 | 100 | 4 | 238 | 3 | 87 | 2 | 15 |
| P56-P63 | BrdU treated mouse 1 | 100 | 1 | 204 | 2 | 78 | 0 | 6 |
| | BrdU treated mouse 2 | 100 | 7 | 236 | 3 | 76 | 0 | 10 |
| | BrdU treated mouse 3 | 100 | 8 | 221 | 2 | 95 | 2 | 24 |
| P63-P70 | BrdU treated mouse 1 | 52 | 5 | 86 | 0 | 40 | 0 | 10 |
| | BrdU treated mouse 2 | 100 | 0 | 206 | 0 | 100 | 1 | 16 |
| | BrdU treated mouse 3 | 72 | 9 | 172 | 2 | 61 | 0 | 11 |

**Third Anagen**

| Treatment period | Sample | Upper ORS | | Lower ORS | | HF Bulb | |
|---|---|---|---|---|---|---|---|
| | | BrdU+ | BrdU- | BrdU+ | BrdU- | BrdU+ | BrdU- |
| P63-P70 | BrdU treated mouse 1 | 6 | 23 | 16 | 39 | 87 | 28 |
| | BrdU treated mouse 2 | 4 | 34 | 46 | 95 | 0 | 0 |
| | BrdU treated mouse 3 | 12 | 37 | 22 | 53 | 103 | 39 |

**Second Anagen**

| Treatment period | Sample | No. of follicles | Upper ORS | | Lower ORS | | HF Bulb | |
|---|---|---|---|---|---|---|---|---|
| | | | BrdU+ | BrdU- | BrdU+ | BrdU- | BrdU+ | BrdU- |
| P21-P30 | BrdU treated mouse 1 | 120 | 75 | 101 | 142 | 136 | 814 | 205 |
| | BrdU treated mouse 2 | 127 | 86 | 99 | 169 | 168 | 486 | 121 |
| | BrdU treated mouse 3 | 128 | 68 | 96 | 147 | 152 | 286 | 63 |

**Supplementary Table 2.** Quantification of BrdU labeling with alternate mouse monoclonal anti-BrdU antibody (clone B44, BD bioscience) during telogen (P49-P56, P56-P63, P63-P70) or anagen (P21-P30). The table lists the total number of HFs counted and total BrdU+ and BrdU - cells from each mouse (n=3). Due to less specificity in staining requiring MOM protocol, BrdU labeling was quantified in vehicle injected mice as well.

**Supplementary Table 2**

#### Second Telogen

| Treatment period | Sample | No. of follicles | SHG | | Bulge | | SG | |
|---|---|---|---|---|---|---|---|---|
| | | | BrdU+ | BrdU- | BrdU+ | BrdU- | BrdU+ | BrdU- |
| P49-P56 | Vehicle treated mouse | 55 | 3 | 208 | 1 | 51 | 0 | 1 |
| | BrdU treated mouse 1 | 116 | 5 | 383 | 0 | 89 | 0 | 10 |
| | BrdU treated mouse 2 | 182 | 4 | 491 | 1 | 141 | 1 | 18 |
| | BrdU treated mouse 3 | 117 | 1 | 349 | 0 | 76 | 0 | 10 |
| P56-P63 | Vehicle treated mouse | 76 | 1 | 199 | 0 | 40 | 0 | 8 |
| | BrdU treated mouse 1 | 204 | 5 | 487 | 2 | 104 | 0 | 10 |
| | BrdU treated mouse 2 | 239 | 7 | 501 | 3 | 137 | 0 | 12 |
| | BrdU treated mouse 3 | 138 | 3 | 281 | 4 | 49 | 0 | 11 |
| P63-P70 | Vehicle treated mouse | 72 | 3 | 159 | 1 | 44 | 0 | 1 |
| | BrdU treated mouse 1 | 130 | 14 | 347 | 2 | 71 | 0 | 2 |
| | BrdU treated mouse 2 | 70 | 3 | 177 | 1 | 44 | 0 | 2 |
| | BrdU treated mouse 3 | 136 | 4 | 279 | 2 | 85 | 0 | 6 |

#### Third Anagen

| Treatment period | Sample | Upper ORS | | Lower ORS | | HF Bulb | |
|---|---|---|---|---|---|---|---|
| | | BrdU+ | BrdU- | BrdU+ | BrdU- | BrdU+ | BrdU- |
| P63-P70 | Vehicle treated mouse | 3 | 92 | 2 | 64 | 2 | 62 |
| | BrdU treated mouse 1 | 35 | 69 | 26 | 64 | 13 | 69 |
| | BrdU treated mouse 2 | 0 | 43 | 0 | 0 | 0 | 0 |
| | BrdU treated mouse 3 | 39 | 112 | 23 | 143 | 9 | 40 |

**Supplementary Table 3.** (a) Quantification of GFP+ melanocytes in control mouse at P30 (n=3). (b) Quantification of GFP+ melanocytes in DOX treated mouse at P30 (n=3). (c) Quantification of GFP retaining melanocytes after DOX administration from P19 to P30 (n=3). (d) Quantification of Ki67 expression in GFP+ melanocytes from P30 DOX treated mice. GFP+ melanocytes from 102 anagen HFs from 3 different DOX treated mice were counted. (e) Quantification of SOX10+ GFP+ and SOX10+ GFP– cells in HFs of *Dct*-H2BGFP mice with and without DOX treatment.

**Supplementary Table 3**

**a**

| Treatment | Sample | No. of follicles | GFP+ melanocytes | | | Total GFP+ cells |
|---|---|---|---|---|---|---|
| | | | Upper ORS | Lower ORS | HF Bulb | |
| P19-P30 Control | Mouse 1 | 100 | 270 | 199 | 1402 | 1871 |
| | Mouse 2 | 100 | 235 | 198 | 1245 | 1678 |
| | Mouse 3 | 100 | 270 | 177 | 1163 | 1610 |

**b**

| Treatment | Sample | No. of follicles | GFP+ melanocytes | | | Total GFP+ cells |
|---|---|---|---|---|---|---|
| | | | Upper ORS | Lower ORS | HF Bulb | |
| P19-P30 DOX | Mouse 1 | 100 | 261 | 231 | 848 | 1340 |
| | Mouse 2 | 100 | 229 | 211 | 1015 | 1455 |
| | Mouse 3 | 100 | 254 | 217 | 838 | 1309 |

**c**

| Treatment period | Sample | No. of follicles | Upper ORS | | Lower ORS | | HF Bulb | |
|---|---|---|---|---|---|---|---|---|
| | | | $GFP^{high}$ | $GFP^{low}$ | $GFP^{high}$ | $GFP^{low}$ | $GFP^{high}$ | $GFP^{low}$ |
| P19-P30 | Mouse 1 | 100 | 141 | 120 | 129 | 102 | 160 | 688 |
| | Mouse 2 | 100 | 145 | 84 | 136 | 75 | 97 | 918 |
| | Mouse 3 | 100 | 181 | 73 | 142 | 75 | 50 | 788 |

**d**

| Sample | Number of Follicles | Upper ORS | | | | Lower ORS | | | | HF bulb | | | |
|---|---|---|---|---|---|---|---|---|---|---|---|---|---|
| | | $GFP^{high}$ | | $GFP^{low}$ | | $GFP^{high}$ | | $GFP^{low}$ | | $GFP^{high}$ | | $GFP^{low}$ | |
| | | Ki67+ | GFP+ | Ki67+ | GFP+ | Ki67+ | GFP+ | Ki67+ | GFP+ | Ki67+ | GFP+ | Ki67+ | GFP+ |
| Mouse 1 | 34 | 19 | 94 | 8 | 29 | 8 | 39 | 3 | 8 | 1 | 5 | 203 | 386 |
| Mouse 2 | 34 | 10 | 74 | 6 | 42 | 10 | 53 | 3 | 13 | 0 | 7 | 161 | 528 |
| Mouse3 | 34 | 12 | 67 | 8 | 37 | 9 | 34 | 4 | 18 | 1 | 5 | 198 | 501 |

**e**

| | Upper ORS | | Lower ORS | | HF bulb | |
|---|---|---|---|---|---|---|
| Sample | GFP+SOX10+ | GFP-SOX10+ | GFP+SOX10+ | GFP-SOX10+ | GFP+SOX10+ | GFP-SOX10+ |
| Control | 36 | 0 | 47 | 0 | 244 | 10 |
| DOX treated | 31 | 5 | 41 | 0 | 296 | 80 |

**Supplementary Table 4.** (a) Cell count for Figures 4 and 5 concerning quantification of high and low expression of markers in $GFP^{high}$ and $GFP^{low}$ melanocytes after doxycycline treatment at P30. The graph lists the markers, HFs counted and the relative expression of the melanogenic markers.

**Supplementary Table 4**

| Marker | Number of Follicles | Upper ORS | | | | Lower ORS | | | | Bulb | | | |
|---|---|---|---|---|---|---|---|---|---|---|---|---|---|
| | | GFP$^{high}$ | | GFP$^{low}$ | | GFP$^{high}$ | | GFP$^{low}$ | | GFP$^{high}$ | | GFP$^{low}$ | |
| | | High | Low | High | Low | High | Low | High | Low | High | Low | High | Low |
| Kit | 40 | 71 | 2 | 26 | 4 | 49 | 4 | 23 | 5 | 3 | 16 | 13 | 399 |
| Nestin | 30 | 40 | 1 | 18 | 0 | 0 | 41 | 0 | 32 | 0 | 4 | 0 | 332 |
| Dct | 31 | 7 | 28 | 2 | 16 | 8 | 21 | 10 | 20 | 10 | 0 | 352 | 4 |
| Tyr | 31 | 6 | 40 | 3 | 13 | 5 | 25 | 7 | 21 | 12 | 0 | 331 | 4 |
| Tyrp1 | 31 | 1 | 38 | 1 | 19 | 2 | 42 | 3 | 38 | 17 | 0 | 355 | 0 |
| Mitf | 50 | 6 | 56 | 4 | 26 | 3 | 69 | 6 | 45 | 24 | 2 | 534 | 17 |
| Pax3 | 31 | 13 | 29 | 5 | 13 | 12 | 24 | 11 | 22 | 6 | 0 | 367 | 38 |
| Ap2α | 30 | 39 | 12 | 14 | 5 | 23 | 8 | 19 | 10 | 10 | 5 | 312 | 47 |

**Supplementary Table 5.** The table lists summary of relative expressions of the markers used to compare the melanocytes in different compartments of the HF after anagen onset doxycycline treatment.

**Supplementary Table 5**

| Marker | Upper ORS | Lower ORS | Bulb |
|---|---|---|---|
| CD34 | high | No expression | No expression |
| Ki67 | low | low | high |
| Kit | high | high | low |
| Nestin | high | No expression | No expression |
| Dct | low | low | high |
| Tyrp1 | low | low | high |
| Tyr | low | low | high |
| Ap-2α | high | high | high |
| Sox10 | high | high | high |
| Pax3 | low | low | high |
| Mitf | low | low | high |

### SUPPLEMENTARY MATERIALS AND METHODS

#### **Immunofluorescence assay with list of antibodies**

Different protocol was followed for mouse monoclonal anti-BrdU antibody (clone B44, BD Bioscience). The sections were first permeabilized with 0.1% Triton-X after HCl treatment for 10 minutes and blocked using streptavidin/biotin blocking kit (vector laboratories) and MOM kit (vector laboratories). Alexa Fluor 546 conjugated streptavidin (1:500 for 15 minutes, Invitrogen) was used as the secondary antibody as per MOM kit protocol.

The sections were incubated overnight in primary antibodies for BrdU (1:200, Rat monoclonal antibody, clone BU1/75 (ICR1), Abcam), Ki67 (1:200, rabbit polyclonal antibody, Leica Biosystems), c-kit (1:200, rat monoclonal antibody, ACK4 clone, Cedarlane), Nestin (1:100, mouse monoclonal antibody, clone rat401, EMD Millipore) Dct (1:200, α-PEP8, rabbit polyclonal antibody, a gift from Dr. Vincent Hearing, NIH), Tyrp1 (1:200, αPEP1, rabbit polyclonal antibody, a gift from Dr. Vincent Hearing, NIH), Tyr (1:200, αPEP7, rabbit polyclonal antibody, a gift from Dr. Vincent Hearing, NIH), Ap-2α (1:100, mouse monoclonal antibody, 3B5 clone, Santa Cruz), Sox10 (1:100, mouse monoclonal antibody, A-2 clone, Santa Cruz), Pax3 (1:200, mouse monoclonal antibody, Developmental Studies Hybridoma Bank) , Mitf (1:100, mouse monoclonal antibody, C5 clone, EMD Millipore), and CD34 (1:200, rat monoclonal antibody, clone Ram34, BD Biosciences). Furthermore, the sections were incubated either in Cy3 conjugated anti-rat or anti-rabbit IgG (H+L) secondary antibody (Jackson Immunoresearch) at 1:1000 dilutions for 1 hour at room temperature or in Alexa Fluor 546 conjugated streptavidin (1:500, for mouse antibodies).

**Distinction of GFP$^{high}$ and GFP$^{low}$ melanocytes in DOX treated mice**

The GFP$^{high}$ and GFP$^{low}$ melanocytes in fluorescence images can be distinguished by difference in fluorescence intensity. The quantitative value for the intensity was calculated as corrected total cell fluorescence (CTCF). The area, integrated density and mean gray value are calculated using Fiji. Background fluorescence is collected adjacent to the cells. The formula used is: CTCF = Integrated Density – (Area of selected cell X Mean fluorescence of background readings). Based on this calculation, we observed that all the GFP$^{low}$ melanocytes are below the CTCF value of 4000 (Figure S3) (Gavet and Pines 2010; McCloy et al. 2014).

**Alkaline phosphatase staining**

The HF cycle stage was confirmed by alkaline phosphatase (AP) staining. A representative section from each of the experiment was first fixed in acetone (stored at -20°C) for 10 minutes and subjected to AP staining following the protocol mentioned by Handjiski et al. using Naphthol-AS-BI-phosphoric acid as substrate and New Fuchsin stock solution (5% in 2N HCl) as coloring component (Handjiski et al. 1994).
